# Accumbal Dopamine and Acetylcholine Dynamics during Psychostimulant Sensitization

**DOI:** 10.1101/2025.03.29.646091

**Authors:** Georg Lange, Federico Gnazzo, Rudolf P. Faust, Jeff A. Beeler

**Affiliations:** Cognitive Neuroscience MS Program, The Graduate Center, City University of New York, New York NY; CUNY Neuroscience Collaborative, Biology and Psychology Programs.; Dept. of Psychology, Queens College, City University of New York.

**Keywords:** Psychostimulant Sensitization, Nucleus Accumbens, Dopamine–Acetylcholine Interactions, D2 Receptors, Cholinergic Interneurons

## Abstract

Behavioral sensitization to repeated psychostimulant exposure is believed to contribute to the development of addiction. Nucleus accumbens (NAcc) dopamine (DA) is known to be a key substrate in sensitization, though recent work suggests that striatal acetylcholine (ACh) may also play a critical role. However, underlying ACh changes and their relationship to DA signaling have not been characterized. Here, we used dual-color fiber photometry to simultaneously measure DA and ACh in the NAcc shell of mice across repeated injections of cocaine or amphetamine. Repeated exposure progressively elevated locomotor activity and increased slow extracellular DA while attenuating transient DA release. Psychostimulants reduced phasic ACh transient amplitude and frequency, an effect that sensitized with repeated injections. However, the temporal coupling of DA and ACh remained unchanged. To determine whether D2 receptors (D2Rs) on cholinergic interneurons (CINs) drive this effect, we generated CIN-selective D2R knockout (KO) mice. Surprisingly, KOs continued to show an acute decrease in ACh and intact DA–ACh correlations after psychostimulant administration. However, they failed to exhibit sensitization of either DA or ACh in response to repeated psychostimulant administration. Despite this lack of sensitization in underlying neuromodulator signaling, the KO mice nevertheless exhibited behavioral sensitization, though at a slower rate than wild-type. These findings suggest that neural sensitization to psychostimulants is dependent on D2R expressed on CINs, but that behavioral sensitization is not dependent on sensitization of these underlying signals.

## 1 Introduction

Psychostimulant sensitization is a process wherein repeated exposure to drugs such as cocaine or amphetamine elicits progressively stronger behavioral responses [1–3]. This sensitization is believed to reflect underlying neural adaptations that mediate the transition from casual drug use to compulsive drug-taking [4–6]. The nucleus accumbens has been the primary focus of investigation and implicated as a crucial substrate for psychostimulant sensitization [7–10]. Psychostimulants increase extracellular accumbal dopamine concentration [8, 11] by blocking or reversing dopamine transporters. This elevated accumbal dopamine facilitates neural plasticity necessary to learn new behaviors and associations that shape future drug responses [12–15]. Early studies using microdialysis confirmed a progressive enhancement of extracellular dopamine on repeated exposures to psychostimulants and that locomotor sensitization is dependent on this accumbal DA [7, 9, 11, 16, 17]. Later work using FSCV to examine DA transmission further confirmed these earlier observations [18, 19].

The primary mechanism of psychostimulants is blockade or reversal of the dopamine transporter (DAT), which increases extracellular dopamine. However, some studies suggest that psychostimulants can affect dopamine signaling through additional mechanisms. Studies using fast-scan cyclic voltammetry (FSCV) found in addition to increased transient decay times that cocaine also increased the amplitude of evoked DA [18–20] and frequency of naturally occurring DA transients [21], leading to hypotheses proposing DAT-independent mechanisms for psychostimulant effects on dopamine [22]. However, more recent fiber photometry experiments challenge those findings [23, 24], with some arguing the previously observed increase in transient DA is an artifact arising from DAT-dependent changes in diffusion [24]. Thus, while the basic mechanism of psychostimulant action is well-established (DAT blockade or reversal), further effects on dopamine signaling remain unclear, and, moreover, whether these additional effects exhibit or contribute to sensitization is poorly understood.

In the NAcc, cholinergic signaling arising from striatal cholinergic interneurons (CINs) interacts with dopamine signaling and could contribute to sensitization. Recently, cholinergic signaling in the striatum has been shown to regulate drug-context learning [25, 26] and extinction [27, 28]. There is a strong reciprocal interaction between mesolimbic dopamine axons and CINs. CINs can depolarize dopamine axons via nicotinic acetylcholine receptors, a circuit strong enough to facilitate dopamine release independent of dopamine cell action potentials [29–32]. Dopamine, on the other hand, modulates CINs via inhibitory dopamine D2 receptors (D2Rs) and excitatory D5 receptors [33–35].

DA and ACh display spontaneous interactions that appear to be governed mainly by extrastriatal inputs rather than local D2R signaling on cholinergic interneurons (CINs) [36]. Yet, other studies indicate that these same D2Rs are essential for psychostimulant sensitization—raising the question of how they shape DA–ACh interactions under repeated drug exposure. For example, selectively knocking out D2Rs in CINs reduces the behavioral response to cocaine and slows locomotor sensitization [37], indicating an important role for CINs for psychostimulant sensitization. Nevertheless, how psychostimulants may alter acetylcholine signaling or any putative reciprocal interactions between DA and ACh in the striatum has not been characterized.

Here, we use dual-color photometry to simultaneously record dopamine and acetylcholine activity in the nucleus accumbens in a repeated injection paradigm of psychostimulant locomotor sensitization using both cocaine and amphetamine. One of the persistent challenges in studying psychostimulant sensitization has been the timescales of measurement tools. Microdialysis captures at best one minute bins, reflecting minute to minute changes, but does not capture DA signaling at the subsecond timescale typically associated with phasic DA activity. FSCV, in contrast, captures phasic DA activity in the form of subsecond DA transients quite well, but because of shifting baseline it can be difficult to use FSCV to examine slower timescales. Here, we endeavor to examine both shortand longtimescales with fiber photometry. We examine DA and acetylcholine separately as well as potential interactions between DA-ACh in a manner similar to Krok et al [36].

We observe locomotor sensitization and progressively greater increases in extracellular dopamine across repeated administrations, consistent with prior literature. At the same time, we observe a concomitant decrease in both the amplitude and frequency of dopamine transients, suggesting differential effects on dopamine signaling at different timescales. In contrast to heightened extracellular dopamine, we also find that psychostimulants diminish acetylcholine signaling. This suppression grows more pronounced with repeated injections, indicating that psychostimulant effects on striatal acetylcholine also sensitize. Notably, we find that these drugs weaken the strength of correlation between DA and ACh on rapid timescales—yet do not alter their relative timing, indicating that psychostimulants reduce DA–ACh coupling without disrupting their fundamental temporal coordination.

We then assessed whether D2Rs on cholinergic interneurons mediate these effects by generating mice selectively lacking D2Rs in CINs. Although this knockout does not affect the acute suppression of ACh transients or the temporal coupling of DA–ACh, it abolishes the progressive sensitization of both DA and ACh that otherwise emerges with repeated drug injections. Moreover, while both wild-type and knockout mice show robust cocaine-induced locomotor sensitization, behavioral sensitization is delayed, occurring more slowly in the knockout mice. Together, these findings indicate that although D2R signaling on CINs is not necessary for the initial drug-induced suppression of ACh or its temporal coordination with DA, it is critical for the progressive sensitization of increased extracelllar dopamine and the parallel suppression of ACh under repeated psychostimulant exposure, though behavioral sensitization remains intact.

## 2 Results

We conducted a psychostimulant sensitization study to investigate how repeated drug administrations of cocaine and amphetamine change locomotor behavior and dopamine (DA) and acetylcholine (ACh) activity in the nucleus accumbens. We measured changes in extracellular DA and ACh concentration with dual-color fiber photometry expressing the GRAB-rDA (dopamine, red) and GRAB-ACh (acetylcholine, green) sensors concurrently in the same mice (Fig. 1a). We recorded infrared videos and trained a DeepLabCut model [38] for locomotor tracking (Fig. 1b). Mice were habituated to an open field chamber after which we recorded an initial baseline where all mice received a saline injection. Then, mice were injected with cocaine (20 mg/kg), amphetamine (2 mg/kg), or saline on five consecutive days with photometry recordings at day 1 and 5. Seven days later, a challenge injection in which all mice, including drug-naive saline controls, received drug was administered and photometry recorded. On each drug administration, mice were placed in the chamber for 10 minutes to establish a pre-injection baseline prior to drug injection and then recorded for an additional 50 minutes post-injection (Fig. 1c). Representative traces from a single animal showing dopamine, acetylcholine and locomotor activity is shown in Fig. 1d.

**Fig. 1:**
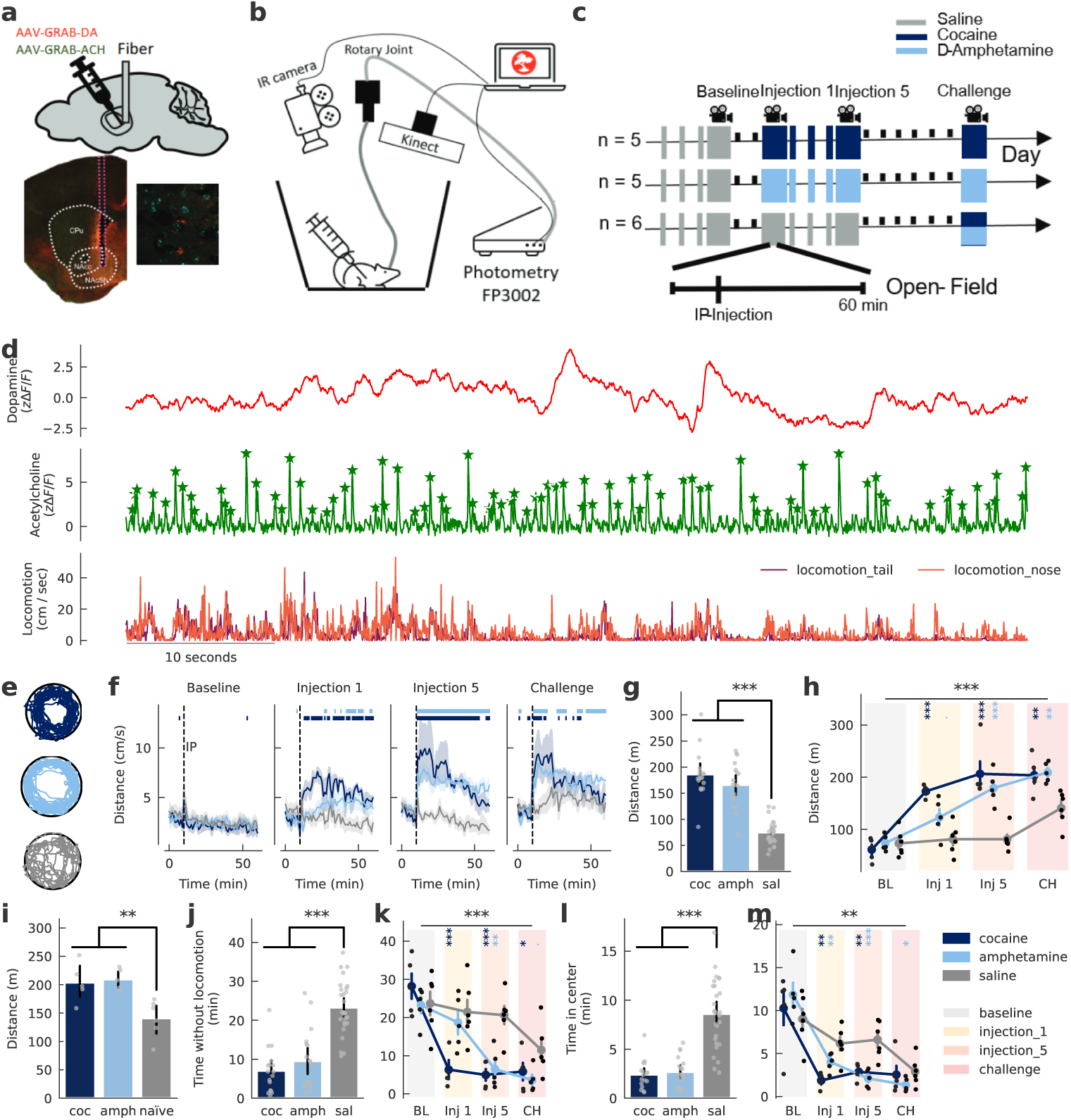
Locomotor sensitization during psychostimulant exposure. **a**, Injection sites of viral GRAB-DA and GRAB-ACh sensors in the left nucleus accumbens (NAcc) shell. Lower left: representative immunohistochemistry (IHC) image showing fiber placement. Lower right: enlarged view of sensor expression (GRAB-DA: green; GRAB-ACh: red; DAPI nuclear staining: blue). **b**, Experimental setup schematic showing simultaneous recording of fiber photometry, infrared, and 3D depth video. **c**, Schedule of psychostimulant sensitization. Tall bars: injection days; thick bars: injection plus recording; short black bars: non-injection days. **d**, Representative traces (1-minute) showing dopamine (DA; top), acetylcholine (ACh; middle), and locomotion (bottom). **e**, Three-minute path traces from representative cocaine-, amphetamine-, and saline-treated animals. **f**, One-minute binned velocity (measured at tail base) across groups and sessions. Horizontal bars above traces indicate significant differences between cocaine (dark blue) or amphetamine (light blue) and saline groups (*p <* 0.05, independent t-test per bin). **g**, Total distance traveled in the 50 minutes post-injection, for each drug condition (Linear Mixed-Effects Model (LME), see Methods 4, *p <* 0.001 for both). **h**, Total distance traveled post-injection by group and session (LME Group*×*Session interaction, *p <* 0.001.) **i**, Distance traveled during challenge (LME, previously exposed cocaine vs. saline, *p* = 0.0025; previously exposed amphetamine vs. saline, *p* = 0.0054). **j**, Total time without locomotion by drug condition (LME, *p <* 0.001 for cocaine and amphetamine). **k**, Resting time by session (LME, Group *×*Session interaction, *p <* 0.001). **l**, Total time in arena center by drug condition (LME, *p <* 0.001). **m**, Arena-center time by session (LME, Group*×*Session interaction, *p* = 0.002). Error bands: s.e.m. (line and point plots) or 95% confidence interval (bar plots). Significance levels:. *p <* 0.1, \**p <* 0.05, \*\**p <* 0.01, \*\*\**p <* 0.001.

### 2.1 Repeated cocaine and amphetamine treatments elevate and sensitize locomotor activity

As expected, psychostimulant administration led to higher locomotor activity than saline controls. This effect was evident in locomotor paths (Fig.1e), elevated velocity (Fig.1f), and greater total distance traveled (Fig.1g). Repeated injections led to progressive enhancement — i.e., sensitization — of these behavioral responses. Average velocity (Fig.1f) and distance traveled (Fig.1h) increased with repeated injections, and mice pre-exposed to psychostimulants traveled longer distances at challenge than drugnaive controls (Fig.1i). Moreover, psychostimulants reduced the time the mouse was immobile (Fig.1j), an effect which appeared to sensitize under amphetamine (Fig. 1k). Additionally, mice treated with cocaine or amphetamine spent less time in the center and more time circling the arena perimeter (Fig. 1e, l–m). Together, these data confirm that both cocaine and amphetamine induce robust locomotor sensitization, evidenced by session-to-session increases and heightened responses at challenge in previously exposed compared to drug-naive mice.

### 2.2 Psychostimulants change DA dynamics across timescales

DA levels fluctuate on different timescales. Established techniques such as microdialysis and FSCV separately capture slower (minutes) and faster (subsecond) signals, respectively. In contrast, fiber photometry (FP) typically captures DA transients in a frequency range from about 0.1 to 15 Hz. When examining slower changes, FP signals can be confounded by low-frequency noise, including bleaching or cable bending. To address this and thereby evaluate longer timescales (for example, the rise and fall of psychostimulant effects over tens of minutes), we developed an artifact-removal method that corrects photobleaching and other artifacts but preserves low-frequency DA fluctuations (Suppl. Fig. 7a, see Methods 3.3).

Using this approach, we are able to resolve DA activity across a wide range of timescales within a single time-series trace for each mouse. In particular, these traces reveal slow fluctuations, rapid phasic transients, and the prolonged DA surge following intraperitoneal (IP) psychostimulant administration (Fig. 2a). While baseline DA appeared stable over time, injections of cocaine or amphetamine led to elevated extracellular DA for at least 40 minutes, with a peak around 5-20 minutes after IP injection (Fig. 2b). This increase was highly reproducible and statistically significant across all mice and sessions (Fig. 2c, d).

**Fig. 2:**
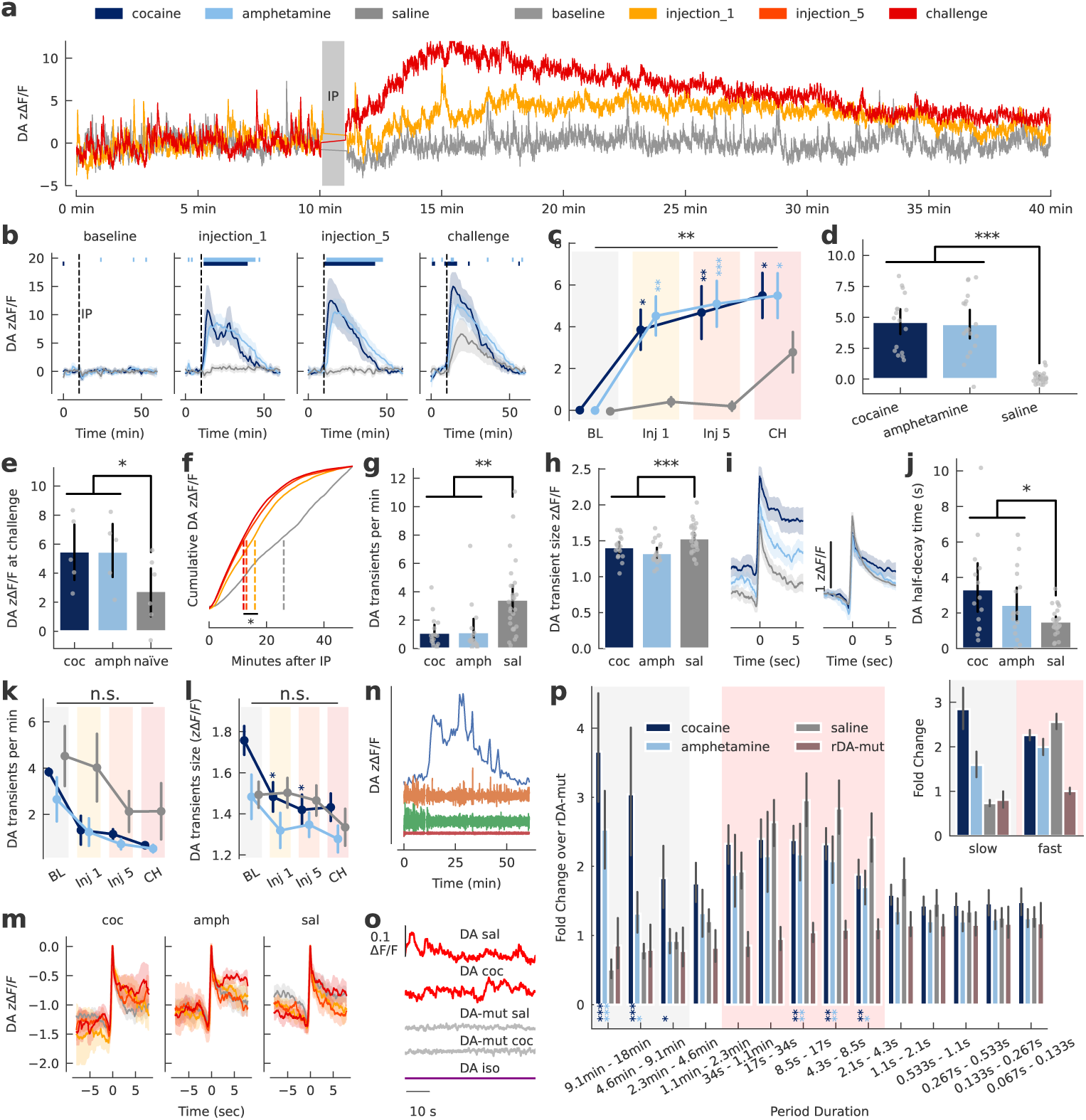
Psychostimulants elevate and sensitize dopamine but attenuate transient signals. **a–d** Psychostimulants increase DA. **a**, Dopamine fluorescence (*z*Δ*F/F* ) from a representative cocaine-treated animal during baseline, injection 1, and challenge sessions. **b**, Mean DA fluorescence (*z*Δ*F/F* ) in 1-minute bins by injection and group. Horizontal bars indicate bins with significant differences between cocaine (dark blue) or amphetamine (light blue) and saline groups (*p <* 0.05, independent t-test). **c**, Mean DA fluorescence post-injection by injection and group (*p* = 0.0093 for Group*×*Session interaction, LME ANOVA). No group differences emerged at baseline (all *p >* 0.96). Injection 1: psychostimulant-treated mice showed increased DA (cocaine: *p* = 0.025; amphetamine: *p* = 0.0032). Injection 5: cocaine (*p* = 0.0015) and amphetamine (*p <* 0.001) exceeded saline. During challenge, saline mice received psychostimulants for the first time, significantly increasing DA (*p* = 0.0024). Previously exposed mice showed additional increases relative to newly treated controls (cocaine: *p* = 0.046; amphetamine: *p* = 0.046). **d**, Mean DA fluorescence post-injection by drug (*p <* 0.001, LME). **e–f** DA sensitization. **e**, DA fluorescence post-injection at challenge (LME, cocaine: *p* = 0.046; amphetamine: *p* = 0.046). **f**, Normalized cumulative DA fluorescence after saline or cocaine injections. Dashed lines indicate when 50% of DA fluorescence is emitted (LME injection 1 vs. challenge, *p* = 0.038). **g–j** Psychostimulants decrease phasic DA transients. **g**, DA transients per minute by drug condition (LME, cocaine: *p <* 0.001; amphetamine: *p* = 0.005 vs. saline). **h**, Amplitude (*z*Δ*F/F* ) of DA transients (LME, *p <* 0.001 both). **i**, DA peri-event traces averaged around the peak of each DA transient (*>* 1 z-score) occurring 0–3 min (left) and 3–50 min (right) post-injection. **j**, Mean half-decay time of DA transients 3–50 min postinjection (LME, cocaine: *p <* 0.001; amphetamine: *p* = 0.017). **k**, Mean DA transients per minute across injections by group (no significant effect of injection). **l**, Mean amplitude (*z*Δ*F/F* ) of DA transients per injection (no significant effect of injection). **m**, DA fluorescence (*z*Δ*F/F* ) normalized and time-locked to the peak of DA transients (first 3 minutes after IP). **n**, Example traces of frequency bands separated by wavelet transforms. **o**, Example traces of GRAB-DA fluorescence (dFF), mut-GRAB-DA fluorescence (DA-insensitive control), and isosbestic signals. **p**, Fold change in DA fluorescence (Δ*F/F* ) relative to mut-GRAB-DA by wavelet frequency band. Bands significantly higher than saline highlighted in grey; significantly higher than mut-GRAB-DA highlighted in pink. The upper right panel shows average fold changes for slow (grey) and fast (pink) frequencies. Error bands: s.e.m. (line and point plots), 95% CI (bar plots). Significance: \**p <* 0.05, \*\**p <* 0.01, \*\*\**p <* 0.001.

Repeated injections increased the magnitude of this elevated DA response indicating psychostimulant-induced sensitization (Fig. 2a-c), also evident in greater responses in previously exposed mice compared to naive (saline) mice at a challenge dose (Fig. 2e). These results align with prior microdialysis studies [7, 9, 11, 16, 17] and suggest that fiber photometry can adequately capture the slower temporal components of DA dynamics. Moreover, we noted a shift in the time at which the DA surge starts, peaks, and decays, which all occurred earlier on subsequent injections (Fig. 2f).

We next tested whether psychostimulants altered the characteristics of phasic DA activity, with a focus on positive transients as a defining feature of phasic release. Cocaine and amphetamine reduced both the frequency (Fig. 2g) and amplitude (Fig. 2h) of DA transients. This decrease was not attributable to group differences or selection of the threshold used for identifying DA transients (Suppl. Fig. 7b). Consequently, the sustained DA surge observed in these experiments does not result from increased frequency of DA release events or greater magnitude of release. Instead, DA gradually accumulates because reuptake is diminished, consistent with the known mechanism of DA transporter blockade. Indeed, transients during the first three minutes after drug administration were comparable in amplitude but decayed so slowly that individual transients no longer return to baseline before the next transient occurs. (Fig. 2i, left). This makes precise half-decay measurements difficult to obtain and we thus only computed half-decay times for DA transients starting 3 minutes after IP, once DA fluorescence had stabilized at a higher level. Transients showed prolonged decay times (Fig. 2i, right), reflected by slower half-decay values (Fig. 2j). These observations are consistent with sustained elevated DA levels emerging from reduced reuptake and not from increased amplitude or frequency of dopamine release..

To assess whether alterations in phasic DA transient activity sensitize, we compared the number and amplitude of DA transients across sessions. Neither effect exhibited a systematic enhancement upon repeated injections (Fig. 2k–l). We next asked if the slowed decay rate of transients sensitizes. Peri-event analyses suggested that under sensitized conditions, DA might remain elevated for longer after each transient (Fig. 2m), though this is difficult to assess quantitatively because, as noted above, basal extracellular DA rises so quickly after injection it is difficult to determine meaningful half-decay times. Despite this uncertainty, our data suggest both the drug-induced DA increase and its enhancement with repeated injections arise from diminished reuptake and not increased amplitude or frequency of dopamine transient events; indeed, these decrease, an effect that apparently does not sensitize with repeated injections.

To characterize DA fluctuations at distinct timescales in a more principled way, we next applied a wavelet transformation to the detrended FP signals. Wavelet transformation decomposes the DA signal into multiple time series, each corresponding to the energy evolution within a specific frequency band over time (Fig. 2n shows example frequency bands). To better estimate the fraction of noise for each frequency band, we additionally recorded mice expressing GRAB-rDA-mut, a version of GRAB-rDA that carries a point-mutation in the sensor’s binding domain that renders it insensitive to DA, thus only capturing random noise (Fig. 2o shows samples of DA signal from each recording condition). We used the GRAB-rDA-mut data to renormalize DA recordings. For mice injected with saline, the wavelet energy in frequency bands of approximately 0.015 Hz (∼1 min period) to 0.46 Hz (∼2 s period) was significantly higher than that of GRAB-rDA-mut controls insensitive to DA (Fig. 2p, frequency bands marked in pink), implying that DA fluctuates spontaneously in this range. Psychostimulant-injected mice showed significantly increased energy in slower frequencies (cocaine: *<* 0.015 Hz; amphetamine: *<* 0.0036 Hz, ∼ 4.6 min period, Fig. 2p, frequency bands marked in grey), which corresponds to the extended DA surge typically lasting tens of minutes. Notably, amphetamine had a more pronounced effect at periods longer than ∼ 4.6 min, while cocaine additionally elevated energy for frequencies with period times of up to ∼ 2.3 min (Fig. 2p, significance indicators below the bars), possibly reflecting their different pharmacokinetics or mechanisms acting on DAT, i.e., cocaine blocking reuptake in an activity-dependent manner while amphetamine additionally reverses transport in an activity-independent manner.

For higher frequency bands, energy is decreased in psychostimulant injected mice compared to saline controls (Fig. 2p, significance indicators below the bars), consistent with the reduced transient activity noted earlier. Overall, both cocaine and amphetamine attenuated transient DA activity, reducing both transient frequency and amplitude, while prolonging DA decay, indicative of substantial reuptake inhibition leading to sustained extracellular DA.

### 2.3 Acetylcholine activity decreases and sensitizes in response to psychostimulants

Acetylcholine (ACh), released by striatal cholinergic interneurons (CINs) in the nucleus accumbens reciprocally regulates DA and can phasically fluctuate in synchrony with DA [36]. However, its role in psychostimulant sensitization remains unclear.

We used GRAB-ACh 3.0 to measure changes in accumbal ACh concentration (Fig. 3a). The signal comprised basal activity interrupted by short (*<* 100 ms), large, positive peaks, i.e., transients, presumably caused by synchronous CIN firing [39] (Fig. 3b). These peaks occurred irregularly at a frequency of approximately 1.3 Hz. Psychostimulants rapidly reduced ACh activity (Fig. 3c) and transient amplitudes declined from about 2.5 z-scores to 1.8 z-scores (Fig. 3d-e). Although they exhibited some recovery, this decrease largely persisted throughout the recording (Fig. 3e) and was significant for both cocaine and amphetamine (Fig. 3f, g).

**Fig. 3:**
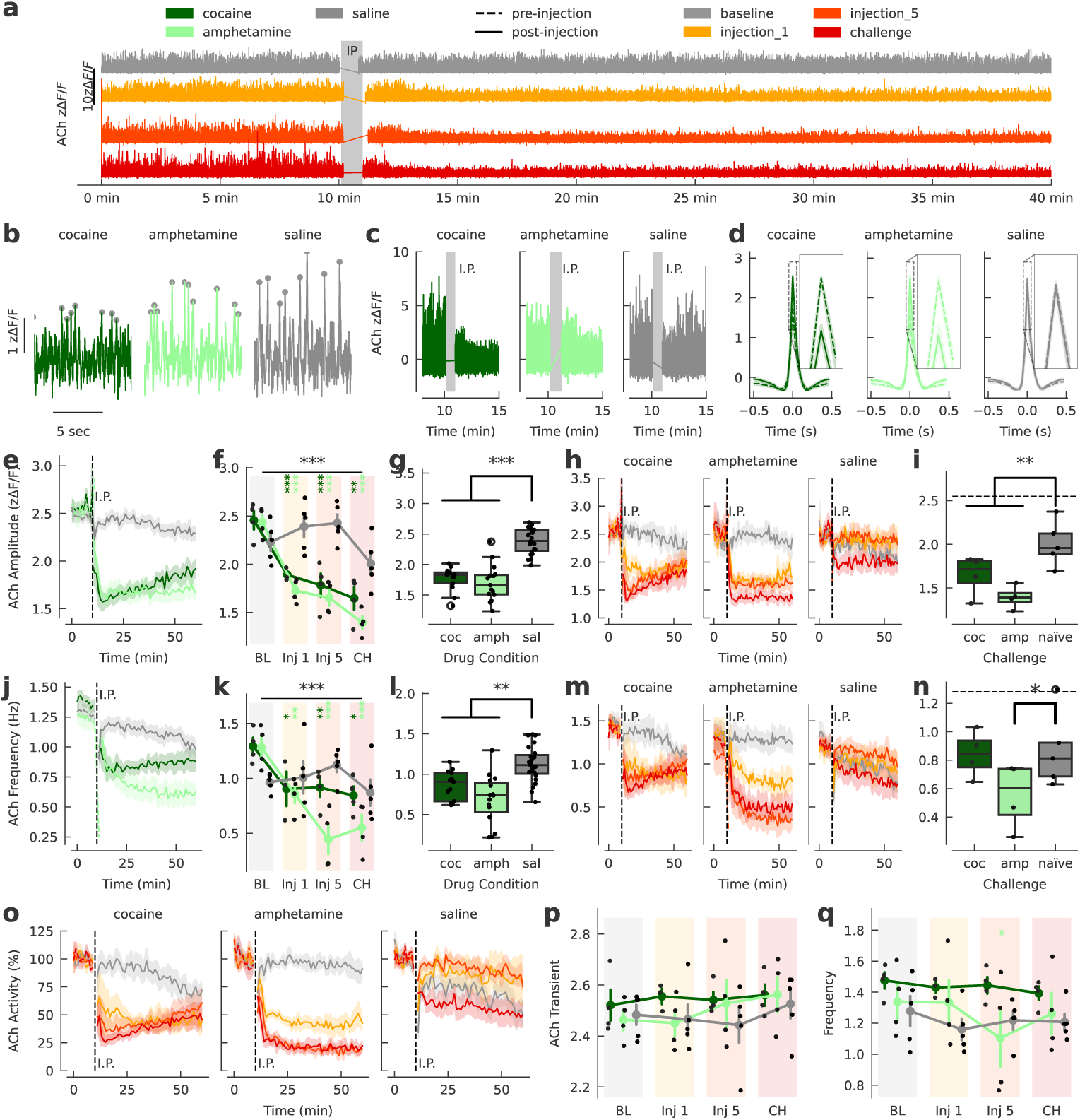
Acetylcholine activity decreases and sensitizes in response to psychostimulants. **a**, ACh fluorescence for each session across injections from a representative animal that received cocaine. **b**, A 10-s example of ACh traces, with identified peaks marked in grey. **c**, ACh traces from representative animals before and after IP injection. **d**, Peri-event plots of ACh fluorescence around ACh transient peaks before (dashed line) and after (solid line) drug injection. **e–i** ACh transient amplitude changes. **e**, Mean amplitude of ACh transients in 1-minute bins. **f**, Mean amplitude of ACh transients post-injection across injections. (LME, *p <* 0.001 for Group*×*Session. LME showed no group differences at baseline but significant interaction effects at injection 1, injection 5, and challenge (*p* = 0.0019 for cocaine*×*challenge; *p <* 0.001 for all other cocaine and amphetamine sessions). **g**, Mean amplitude of post-injection ACh transients by drug condition (LME, *p <* 0.001). **h**, Mean amplitude of ACh transients in 1-minute bins with repeated injections and challenge superimposed for each group. **i**, Mean amplitude of ACh transients after challenge injection of cocaine or amphetamine. Coc and amph indicates animals previously exposed to these in injections 1-5. The dashed line denotes the mean amplitude averaged across groups during the 10-minute baseline (LME, cocaine: *p* = 0.0019 or amphetamine: *p <* 0.001 vs. drug-naive mice). **j-n** ACh transient frequency changes. **j**, Mean frequency of ACh transients in 1-minute bins. **k**, Mean frequency of ACh transients post-injection across injections (*p <* 0.001 for Group*×*Session interaction, LME ANOVA). Cocaine-treated mice showed reduced frequency at injection 1 (*p* = 0.016), 5 (*p* = 0.0022), and challenge (*p* = 0.034); Amphetamine-treated mice showed reduced frequency at injection 1 (*p* = 0.0062), 5 (*p <* 0.001), and challenge (*p <* 0.001). **l**, Mean frequency of post-injection ACh transients by drug condition (LME, *p* = 0.0034). **m**, Mean frequency of ACh transients in 1-minute bins for repeated injections and challenge superimposed for each group. **n**, Mean frequency of ACh transients after challenge injection of cocaine or amphetamine. Coc and amph indicates animals previously exposed to these in injections 1-5. The dashed line denotes the mean transient frequency averaged across groups during the 10-minute baseline (LME, amphetamine: *p* = 0.021; cocaine: *p* = 0.84, n.s.). **o**, Mean ACh activity in 1-minute bins, activity computed as the sum of ACh transient amplitudes in each bin, normalized to the mean activity of the 10-minute pre-injection baseline (for each recording). **p**, Mean ACh transient amplitude during the 10-minute pre-injection baseline across sessions by group. No significant differences observed. **q**, Mean ACh transient frequency during the 10-minute pre-injection baseline across sessions by group. No significant differences observed. Error bands: s.e.m. (line and point plots), 95% CI (bar plots). Significance: \**p <* 0.05, \*\**p <* 0.01, \*\*\**p <* 0.001.

Repeated injections enhanced this reduction in ACh transient amplitudes (Fig. 3h) and mice previously exposed to drug showed a greater reduction than naive (saline control) mice at challenge injection (Fig. 3i), indicating sensitization of this psychostimulant-induced reduction in ACh transient amplitude. Notably, cocaine sensitization was most evident over the first 15 minutes but with gradual recovery that converged to a stable point across injections whereas amphetamine sensitization remained evident for the entire duration of the recording (Fig. 3h), again likely reflecting differences in mechanism of action at DAT.

We repeated this analysis to examine the frequency of ACh transients. Transient frequency remained stable after saline injections, but after psychostimulant administration declined from 1.3 Hz to approximately 0.86 Hz for cocaine and from 1.3 Hz to 0.69 Hz for amphetamine, remaining at these reduced levels throughout the recording (Fig. 3j). This effect appears to sensitize in response to repeated amphetamine but not cocaine injections (Fig. 3k). Despite this difference in sensitization, the overall reduction in frequency is significant for both amphetamine and cocaine (Fig. 3l). Looking at time course of reduced transient frequency post-drug injection, amphetamine induces sensitization that remains throughout the recording session (Fig. 3m). Consistent with this observation, on challenge dose only mice previously exposed to amphetamine but not cocaine showed a greater decrease in Ach transient frequency compared to drug-naive (saline control) mice (Fig. 3n), again likely reflecting different underlying mechanisms of action at the DAT receptor.

To obtain a more principled measure of the combined effect of decreased transient amplitude and frequency on Ach activity and ’tone,’ we combined both parameters into a single index by summing the amplitudes of all transients within 1-minute bins and normalizing them to the mean of pre-injection bins, i.e., for each session the 10minute pre-injection baseline average of these bins was set to 100%. We refer to this normalized quantity as “ACh activity,” which serves as an overall indicator of ACh release. As sensitization of effects on Ach transients is different between the drugs, it is unsurprising that this activity measure sensitized in a similar manner: cocaine induced a rapid sensitized drop in activity that then gradually recovered, whereas the sensitized amphetamine induced reduction of Ach activity remained persistent throughout the recording (Fig. 3o). After the first injection, activity typically fell by about 50%, but in sensitized mice it decreased by as much as 75-80% (Fig. 3o). As above, we see gradual, partial recovery in cocaine treated mice where sensitization is reflected largely in the initial response, while sensitization of the more robust effects observed in amphetamine treated mice are maintained across the entire duration of recording.

Finally, it appears possible that psychostimulants not only cause acute changes post-injection but might modify baseline ACh activity in subsequent sessions (Fig. 3a). To test this, we examined whether psychostimulant exposure led to changes during the 10 minute pre-injection period. We compared the amplitude and frequency of ACh transients during the 10-minute pre-injection period across groups and injections but found no significant differences (Fig. 3p–q).

### 2.4 Psychostimulants reduce transient DA–ACh correlations without altering their temporal alignment

Here, we examine the relationship between DA and ACh. Under baseline conditions, DA and ACh in the nucleus accumbens display distinct dynamic profiles: DA fluctuates slowly (over many seconds), while ACh is dominated by rapid transients (Fig. 4a). After psychostimulant injection, basal DA is greatly increased, though frequency and amplitude of transients are diminished, while ACh activity is blunted (Fig. 4b, note scale change on right axis). At baseline and after saline injection, cross-correlation analysis revealed pronounced positive and negative correlations, indicating robust coordinated fluctuations between DA and ACh (Fig.4c). On average, the strongest negative cross-correlation was at −170 ms and the maximum positive at 320 ms relative to DA (Fig. 4d). Cocaine or amphetamine injection significantly reduced both positive and negative correlation peaks (Fig.4e,g). This effect does not appear to sensitize (Fig. 4f,h). At the challenge session, mice previously exposed to cocaine showed weaker positive but not negative peak correlations compared to both drug-naive controls and mice previously exposed to amphetamine; however, this likely reflects a more robust decrease in max positive correlation in cocaine overall rather than compelling evidence for sensitization.

**Fig. 4:**
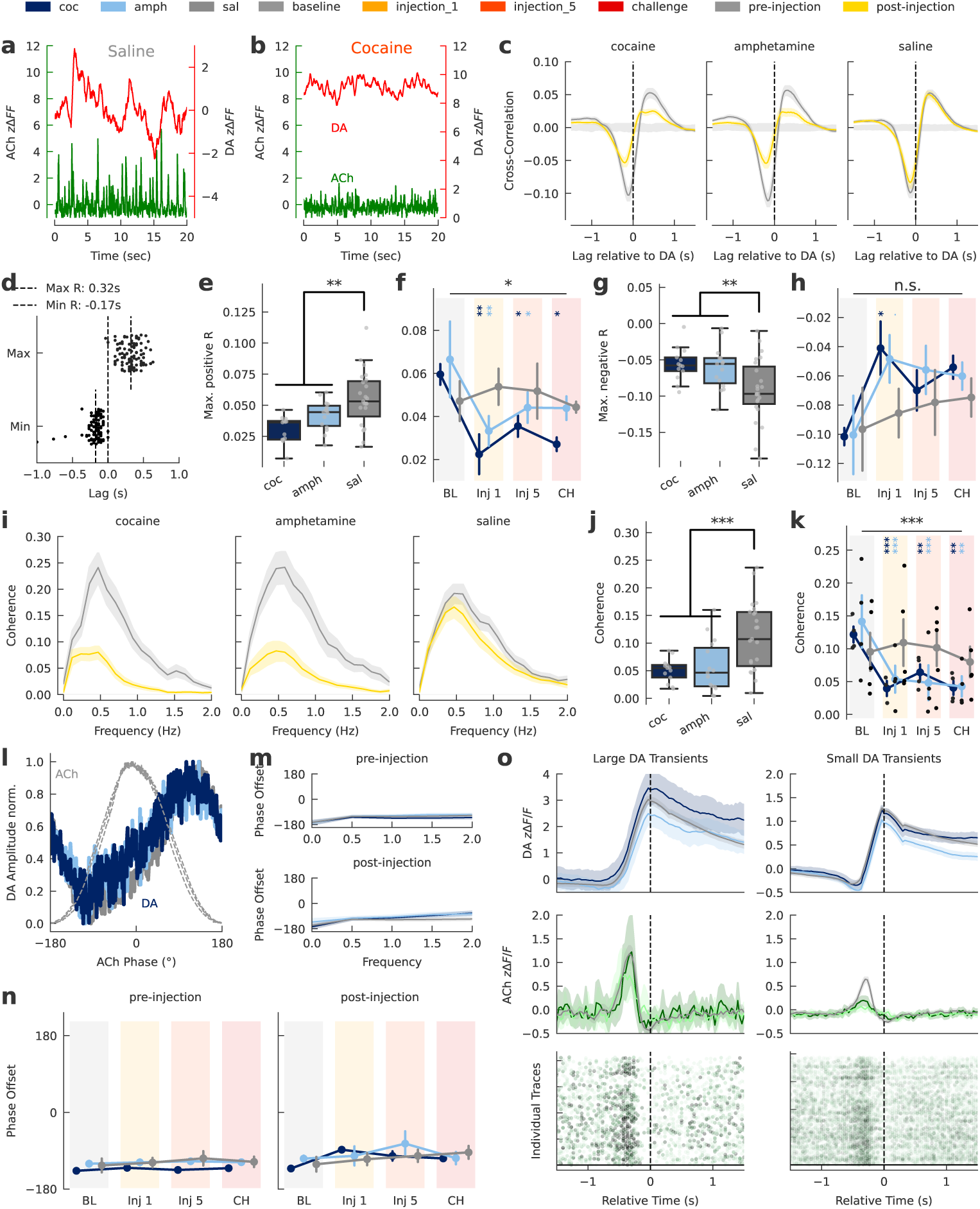
Psychostimulants reduce transient DA–ACh correlations without altering their temporal alignment. **a**, A 20-s DA (red) and ACh (green) trace from a representative animal. **b**, DA and ACh from the same animal shown in **a** after cocaine administration. **c**, Cross-correlation between DA and ACh before and after IP injection. **d**, Time lag (s) of the largest positive and negative cross-correlation post-injection, each dot represents the average of one recording. **e**, Max. positive Pearson’s R between DA and ACh fluorescence by drug condition (LME, *p <* 0.001 cocaine, *p* = 0.004 amphetamine vs saline). **f**, Max. positive Pearson’s R by group and session (LME, *p* = 0.030 for Group*×*Session interaction; cocaine *×* injection 1: *p* = 0.002; amphetamine*×*injection 1: *p* = 0.0035; cocaine *×* injection 5: *p* = 0.0298; amphetamine*×*injection 5: *p* = 0.0398; cocaine *×* challenge: *p* = 0.025; amphetamine*×*challenge: *p* = 0.12). **g**, Max. negative Pearson’s R between DA and ACh fluorescence by drug condition (LME, *p <* 0.001 cocaine, *p* = 0.006 amphetamine vs saline). **h**, Max. negative Pearson’s R by group and session (LME, Group*×*Session interaction: *p* = 0.42). **i**, Coherence by drug administered postinjection. **j**, Mean coherence (0–1 Hz) between DA and ACh fluorescence by drug condition (LME, *p <* 0.001). **k**, Mean coherence (0–1 Hz) by group and session (LME, *p <* 0.001 for Group*×*Session interaction; cocaine *×* injection 1: *p <* 0.001; amphetamine*×*injection 1: *p <* 0.001; cocaine *×* injection 5: *p* = 0.0091; amphetamine*×*injection 5: *p <* 0.001; cocaine *×* challenge: *p* = 0.0074; amphetamine*×*challenge: *p* = 0.001). **l**, Mean normalized DA fluorescence versus phase of periodic ACh fluctuations (grey for reference). **m**, Phase offset of DA versus ACh fluorescence by drug administered. **n**, Phase offset (degrees) between DA and ACh pre-injection (LME, no significant effects) and post-injection (LME, no significant effects). **o**, Upper panel: DA fluorescence time-locked to peaks of detected DA transients by drug administered. Middle panel: mean ACh fluorescence during the same events. Bottom panel: individually detected ACh events around DA transients, colored by height (dashed lines separate animals). Left panels show DA transients larger than 2 z-scores; right panels show transients between 1–2 z-scores. Error bands: s.e.m. (line and point plots), 95% CI (bar plots). Significance: \**p <* 0.05, \*\**p <* 0.01, \*\*\**p <* 0.001.

We next examined frequency-specific interactions using coherence analysis. Initially, DA and ACh signals were most coherent at 0.5–1 Hz. After psychostimulant treatment, coherence in this band significantly decreased (Fig.4i-k), indicating that the coordination at this key frequency range is selectively disrupted. At frequencies outside this band, coherence was nearly zero, leaving little room for alterations in response to psychostimulants or analysis. Despite these changes in amplitude-based correlation-coherence, the relative *timing* between DA and ACh signals remained stable. The phase offset, approximately 160–180° at baseline, did not shift after psychostimulant exposure (Fig. 4l-n). Thus, drugs reduce the strength of DA–ACh coupling without altering their fundamental temporal alignment.

Event-triggered analyses, where we grouped transients by size, revealed that although the overall amplitude and frequency of DA and ACh transients are reduced by psychostimulants (as described above), the timing relationship between them remains consistent. In all groups, a DA transient of a given magnitude was reliably preceded by an ACh transient of comparable relative size (Fig. 4o). Further, we found that the size of DA transients significantly predict the size of the preceding ACh transient (LME, *p <* 0.001, a 1-unit DA increase predicts a 0.53-unit increase in ACh transient amplitude) and this relationship is not altered for cocaine (LME, *p* = 0.47) or amphetamine (LME, *p* = 0.36).

Overall, these results indicate that psychostimulants diminish the strength of the coordinated relationship between DA and ACh signaling — as evidenced by reduced cross-correlation and coherence — while preserving their temporal alignment. This dissociation between a relationship in amplitude modulation between the signals and temporal coordination may be important to understanding the interactions between dopamine and acetylcholine in the striatum.

### 2.5 D2R knockout in cholinergic interneurons delays behavioral sensitization

After characterizing psychostimulant-induced changes of accumbal ACh activity, we asked whether disrupting DA-ACh interaction alters drug responses and sensitization. Specifically, we deleted dopamine D2 receptor (D2R) in striatal CINs by crossing mice expressing cre-recombinase under control of the choline-acetyltransferase promoter (chat-cre mice) with a floxed D2R line to selectively knock out D2R from cholinergic cells. Knockouts (KO) and wild-type (WT) littermates underwent the same experimental protocol as above. As there were no saline controls in this study, the ’challenge’ dose is simply a final (6th) test injection a week following the repeated injection protocol.

We observed no locomotor differences between KO and WT littermates during the baseline recording (Fig. 2.4a, left). After the first cocaine injection, KO mice displayed significantly less locomotor activity compared to WT controls (Fig. 2.4b, c), suggesting that D2R on CINs mediate a component of the initial, naive response to cocaine. However, with repeated injections, both KO and WT mice exhibit locomotor sensitization, though sensitization appears slower in KO mice (Fig. 2.4b, d, e). At challenge, there is no difference in locomotor response between the genotypes (Fig. 2.4a, b, d). A prior study using the same D2R CIN-KO mice [37] also observed delayed locomotor sensitization. In that study however, the effect was more pronounced and prolonged. This difference in timecourse likely arises from differences in dose (10 mg/kg in the Lewis et al study vs. 20 mg/kg here), suggesting this delayed sensitization in CIN D2R KOs is dose-dependent. Both studies, however, demonstrate that D2R on CINs accelerate but are not required for sensitization to psychostimulants. Similarly, KO mice compared to WT spent slightly more time without locomotion after cocaine injection 1 (*p* = 0.07; Fig.2.4f) but showed no difference in center time (Fig. 2.4g). In summary, removing D2Rs from CINs moderately blunts the initial locomotor response to cocaine and slows the onset of sensitization, both effects likely dose-dependent.

### 2.6 D2R knockout in CINs prevents DA sensitization

We examined whether deletion of D2R altered underlying dopamine signaling in response to psychostimulants. KOs show greater inter-individual variability in dopamine response to psychostimulants and mostly did not show sensitization of this dopamine response, while WT exhibited more consistency across mice and all exhibited sensitization (Fig. 2.4h). These trends are evident when aggregate data across injections is plotted separately for each genotype (Fig. 2.4i, colored bars at top indicate timepoints of significant difference between injection 5 or challenge and injection 1). Zeroing in on the first 10 minutes post-injection (Fig. 2.4j), we again observe greater variability and a lack of DA sensitization of the DA repsonse in KO mice. As expected, WT mice clearly exhibited sensitization in the dopamine response at both injection 5 and the challenge injection (compared within animal to injection 1) while the KO mice did not (Fig. 2.4k-l). Thus, deletion of D2R in CINs prevents sensitization of increased DA in response to repeated psychostimulant administration.

### 2.7 CIN-Specific D2R Deletion Abolishes Cocaine-Induced ACh Sensitization

We next investigated how deleting D2Rs from CINs affects the progressive reduction of ACh activity under repeated cocaine injections (Fig. 6a-c). In both WT and KO mice, ACh transient amplitudes decreased after cocaine (Fig. 6d-e), with no genotype differences after the first injection (Fig. 6f), indicating that the acute reduction of ACh transient amplitude in response to psychostimulants is not mediated by CIN D2Rs. We asked whether this decrease in amplitude of ACh transients sensitizes in KO as in WT mice (§2.3). WT mice exhibited sensitization while KO mice did not (Fig. 6g-i), suggesting that deleting CIN D2Rs abolishes cocaine-induced ACh sensitization.

**Fig. 5:**
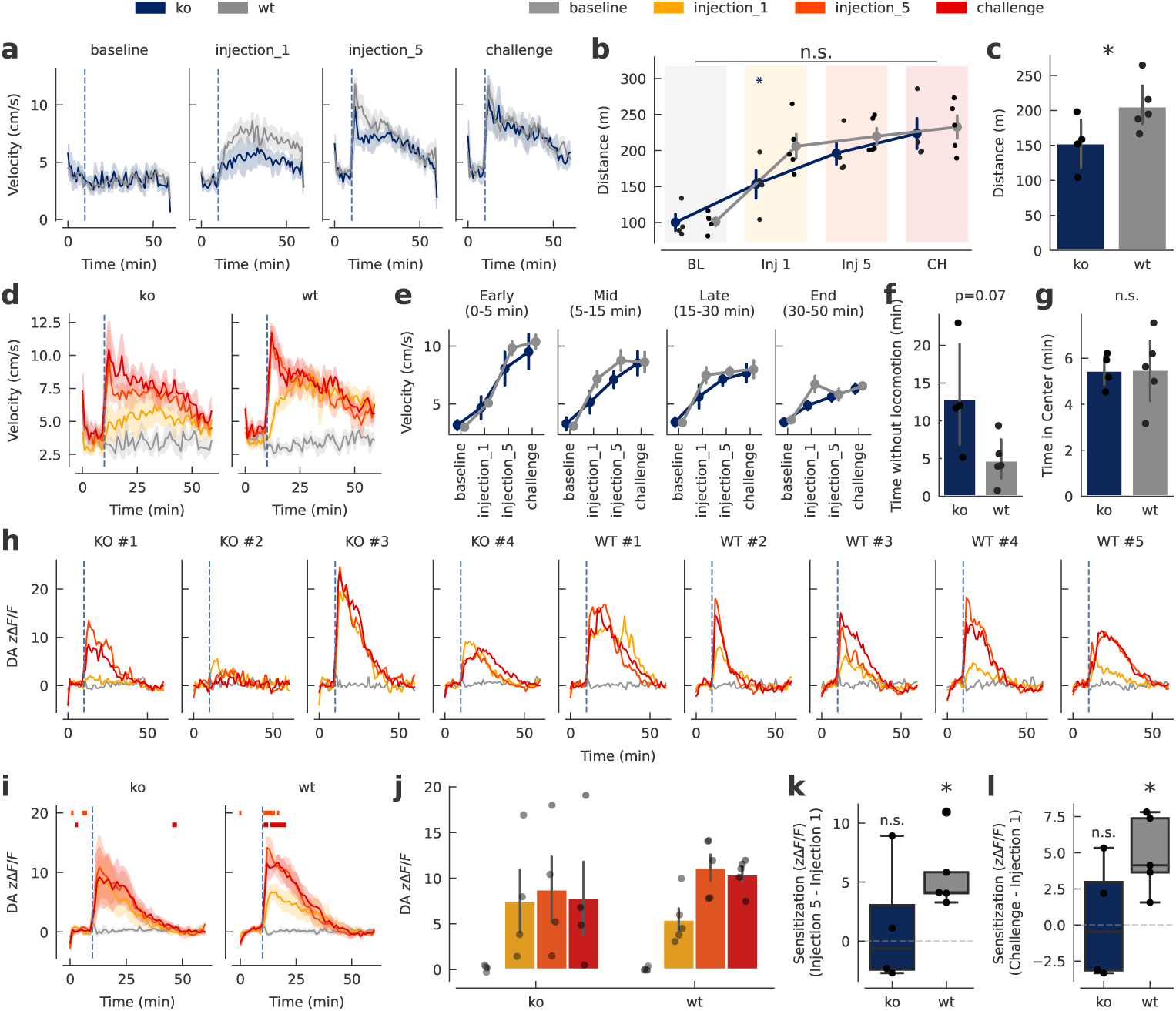
Conditional D2R Knockout in CINs prevents cocaine-evoked DA sensitization. **a**, Mean velocity (1-minute bins) measured at the base of the tail using DeepLabCut.. Dashed line marks the time of injection (10 min). **b**, Total distance traveled (m) post-injection (LME, drug vs. saline, *p <* 0.001). **c**, Total distance traveled after the first cocaine injection, (LME, *p* = 0.017). **d**, Time course of locomotor velocity, dashed line marks the injection time. **e**, Locomotor responses divided into early (0–5 min), mid (5–15 min), late (15–30 min), and end (30–50 min) post-injection phases. **f**, Total time without locomotor activity (LME, *p* = 0.073). **g**, Total time spent in the center. **h**, Mean (1-minute bins) standardized DA fluorescence (*z*Δ*F/F* ) for each animal across sessions. **i**, Time course of DA activity (*z*Δ*F/F* ) across sessions for each genotype. Colored bars above indicate significant increases at injection 5 or challenge relative to the first cocaine injection. **j**, Mean dopamine fluorescence during the first 10 min post-injection. **k**, Within-animal DA sensitization at injection 5, calculated as the difference in mean DA zdFF (0–10 min post-injection) between injection 1 and 5. Wild-type mice showed significant sensitization (*p* = 0.015, one-sample ttest against zero), whereas knockout mice did not (*p* = 0.68). **l**, Within-animal DA sensitization at challenge, computed as the difference in mean DA zdFF (0–10 min post-injection) between injection 1 and challenge. Wild-type mice showed significant sensitization (*p* = 0.014), whereas knockout mice did not (*p* = 0.90). Error bands: s.e.m. (line and point plots), 95% CI (bar plots). Significance: \**p <* 0.05, \*\**p <* 0.01, \*\*\**p <* 0.001.

**Fig. 6:**
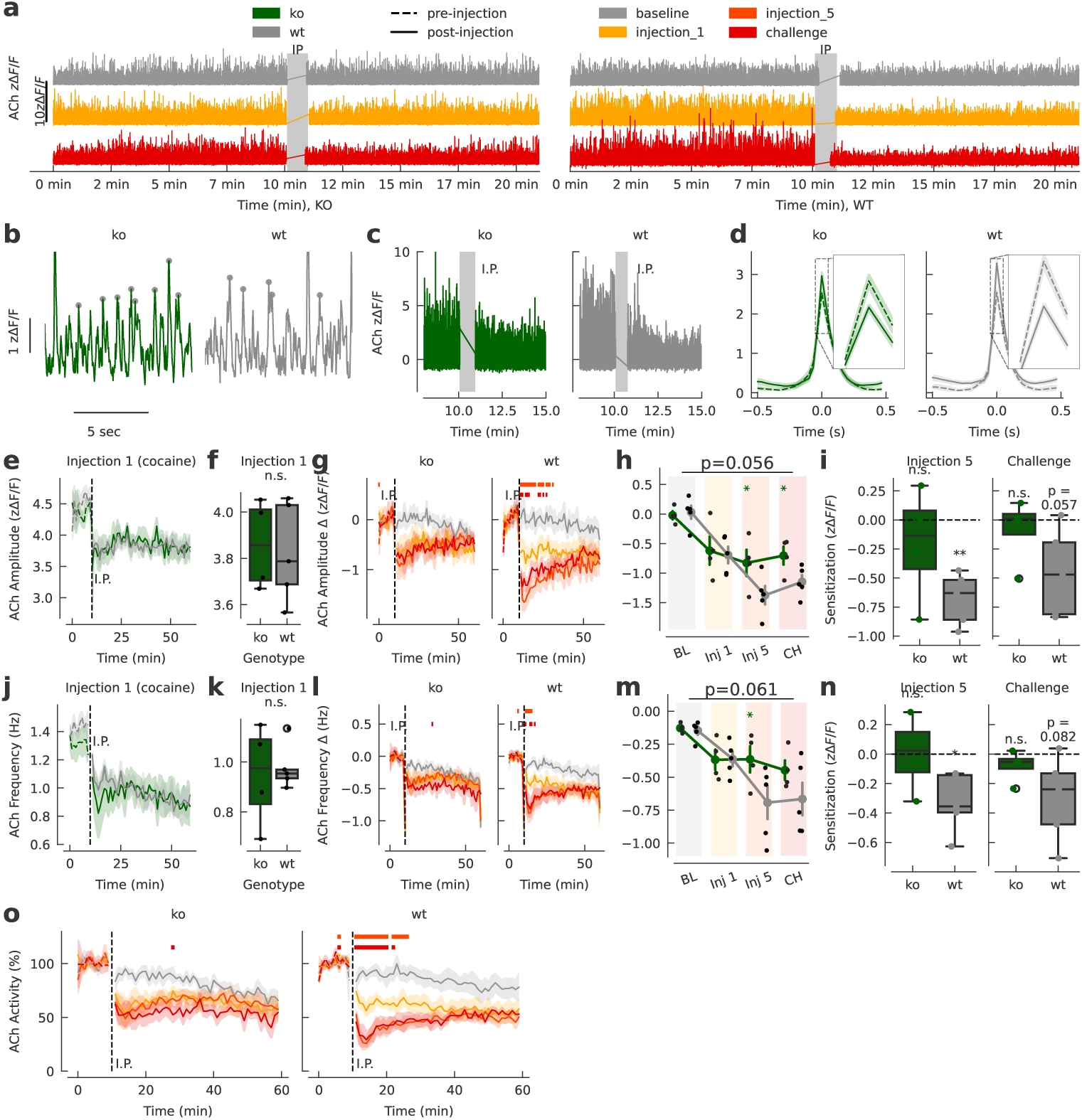
CIN-Specific D2R Deletion Abolishes Cocaine-Induced ACh Sensitization. **a**, ACh fluorescence around the time of IP injection for an example KO (left) and WT (right) mouse during baseline (grey), the first cocaine injection (yellow) and challenge (red). **b**, A 10-s example of ACh traces, with identified peaks marked in grey. **c**, ACh traces from representative animals around the time of IP injection. **d**, Peri-event plots of ACh fluorescence around ACh events before (dashed line) and after (solid line) cocaine injection. **e**, Mean amplitude (*z*Δ*F/F* ) of ACh transients in 1-minute bins during the first cocaine injection. **f**, Mean amplitude of ACh transients after the first cocaine injection. **g**, Mean change in ACh transient amplitude compared to baseline in 1-minute bins. The baseline was calculated for each recording individually as the mean ACh transient amplitude during the 10-minute preinjection period. Colored bars indicate bins where injection 5 (orange) or challenge (red) data were significantly lower than injection 1. **h**, Mean change in ACh transient amplitude post-injection. (LME: No significant Genotype differences at baseline and injection 1. *Genotype x Session* interaction: *p* = 0.056; *Genotype x injection 5* (*p* = 0.019); *Genotype x challenge* (*p* = 0.048). Moreover, there was a session main effect for injection 1, 5, and challenge (*p <* 0.001). **i**, Within-animal ACh sensitization at injection 5 (left panel) and challenge (right panel), calculated as the difference in mean ACh transient amplitude (0–10 min post-injection) between injection 1 and 5 or challenge. Wild-type mice showed significant sensitization (injection 5: *p* = 0.023; challenge: *p* = 0.082 one-sample t-test against zero), whereas knockout mice did not (injection 5: *p* = 0.98; challenge: *p* = 0.24). **j**, Mean frequency (Hz) of ACh transients in 1-minute bins during the first cocaine injection. **k**, Mean frequency of ACh transients after the first cocaine injection. **l**, Mean change in ACh transient frequency compared to baseline in 1-minute bins. The baseline was calculated for each recording individually as the mean ACh transient frequency during the 10-minute preinjection period. Colored bars indicate bins where injection 5 (orange) or challenge (red) data were significantly lower than injection 1. **m**, Mean change in ACh transient frequency post-injection. (LME: no significant Genotype differences at baseline and injection 1, *Genotype x Session* interaction: *p* = 0.061; *Genotype x Injection 5* : *p* = 0.031; *Genotype x Challenge*: *p* = 0.15). **n**, Within-animal ACh sensitization at injection 5 (left panel) and challenge (right panel), calculated as the difference in mean ACh transient frequency (0–10 min post-injection) between injection 1 and 5. Wild-type mice showed significant sensitization (injection 5: *p* = 0.002; challenge: *p* = 0.057 one-sample t-test against zero), whereas knockout mice did not (injection 5: *p* = 0.46; challenge: *p* = 0.60). **o**, Mean ACh activity per genotype and session in 1-minute bins, defined as the sum of ACh transient amplitudes in each bin, normalized to the 10-minute pre-injection baseline. Error bands: s.e.m. (line and point plots), 95% CI (bar plots). Significance: \**p <* 0.05, \*\**p <* 0.01, \*\*\**p <* 0.001.

We similarly examined changes in the frequency of ACh transients. Both WT and KO mice showed a marked decrease in frequency of ACh transients after cocaine injection, again with no genotype difference observed at injection 1, though we observe greater variability in the KO mice (Fig. 6j,k). In WT mice, however, this reduction of ACh transient frequency in response to cocaine sensitized across repeated injections while it remained at the level of injection 1 for KO mice (Fig. 6l-n).

Finally, we combined amplitude and frequency into a single ACh activity index as above (§2.3). This combined measure decreased by about 45% following the first cocaine injection for both genotypes (Fig. 6o). The WT mice show further decreases across subsequent injections, indicating sensitization, whereas KO mice do not exhibit progressively greater reductions in ACh activity (Fig. 6o). In sum, removing D2Rs from CINs abolishes the progressive reduction of ACh activity observed with repeated cocaine exposure, indicating that CIN D2Rs are essential for ACh sensitization.

### 2.8 Conditional D2R KO Does Not Alter DA–ACh Coordination

We next asked whether the KO alters interactions between DA and ACh. Despite marked differences in DA and ACh sensitization, the transient-level and frequency-domain coordination of DA and ACh appeared unaffected by genotype (Suppl. Fig.8 a, b). As in earlier experiments, cocaine significantly reduced the positive crosscorrelation between DA and ACh in both WT and KO mice (Suppl. Fig. 8c, d), with no additional KO effect on the magnitude or time course of the cross-correlation (Suppl. Fig.8d–h). Cocaine broadly decreased coherence (Suppl. Fig.8i) but the KO had no effect on baseline coherence (Suppl. Fig.8j) or changes over sessions (Suppl. Fig.8k). Similarly, the phase alignment of DA and ACh signals also remained same across genotypes (Suppl. Fig. 8l–n).

Event-triggered analyses supported this conclusion: KO mice exhibited the same DA-triggered ACh patterns as WT mice, showing no differences in amplitude or timing of ACh peaks surrounding DA transients (Suppl. Fig. 8o). Further, we found that the size of DA transients significantly predict the size of the preceding ACh transient (LME, *p <* 0.001) and this relationship is not altered for KO (LME, *p* = 0.71).

Thus, while D2R deletion in CINs disrupts DA and ACh sensitization, it does not measurably alter phasic DA–ACh coordination at either the transient or coherence level.

## 3 Discussion

Cocaine and amphetamine elevate extracellular DA by blocking the dopamine transporter (DAT), as shown by microdialysis [40–42], fast-scan cyclic voltammetry (FSCV) [19–21, 43], and more recently fiber photometry [23, 24]. Prior studies using FSCV additionally suggested that cocaine also increases phasic activity observed as increased frequency and amplitude of DA transients [21, 43]. In contrast, newer FP data [23, 24] and re-analyses of FSCV measurements suggest these increases in transient activity might be artifactual: DA diffusion around the electrode is altered when DAT is blocked and leads to measurement errors [24]. Here, we provide additional photometry evidence indicating that psychostimulants attenuate phasic DA activity, decreasing both the amplitude and frequency of DA transients. Prior studies using both FSCV [21] and photometry [44] have demonstrated a critical role for D2 autoreceptors in regulating DA neuron activity in response to psychostimulants, regulating both dopamine cell activity in the midbrain (frequency) and release at dopamine terminals (amplitude, though amplitude also depends upon dopamine cell synchrony [45]). This suggests a straightforward interpretation: as psychostimulants induce greater accumulating extracellular dopamine, D2 autoreceptor activation increases providing greater feedback inhibition, decreasing both dopamine cell activity and terminal release, thus reducing both the frequency and amplitude of dopamine transients. Extracellular dopamine is effectively the averaging of release events. The temporal window across which release events are averaged is controlled by the rates DAT reuptake [46]. By blocking DAT, psychostimulants effectively change this averaging function, increasing the width of the time averaging window and effectively amplifying slower timecourse, averaged activity. Our data suggest more rapid, phasic signaling– widely believed to encode more temporally precise, behaviorally salient information that mediates decision-making and behavior [47] – is diminished. This suggest that psychostimulants alter the relationship between slow and fast timecourse DA signaling, an effect that may be central to understanding their cognitive-behavioral effects.

### 3.1 Acetylcholine, psychostimulants and sensitization

We found that psychostimulants rapidly decrease acetylcholine activity, an effect that sensitizes. Early studies using microdialysis or cFOS activation suggested that psychostimulants increase striatal acetylcholine [48–50] and CIN activity [51]. Cocaine applied to nucleus accumbens slices increases CIN activity [35]. In that same study, Witten et al [35] demonstrated that optogenetically inhibiting CIN activity diminishes cocaine conditioned place preference (cCPP), though increasing CIN activity does not conversely enhance cCPP. In contrast, Guzman et al [52] genetically deleted the acetylcholine vesicular transporter from striatal cholinergic interneurons, selectively eliminating acetylcholine but not glutamate release from these cells. Despite numerous studies suggesting CINs are critical to psychostimulant response [25–28], ablating striatal cholinergic signaling had surprisingly limited effects, with only modest dosedependent effects on locomotor activity, mildly decreased at the mid-dose range (20 mg/kg, as we used here) with no effect at low or high doses. In contrast to the Witten study [35], removing cholinergic signaling from CINs altogether did not alter cCPP; however, it did enhance locomotor sensitization. Finally, Ztaou et al [53] comprehensively assessed CIN activation in response to 2 or 16 mg/kg of amphetamine using phosphorylation of ribosomal protein S6 at serine 240 and 244 residues (a marker of CIN activity, [54]). At the lower (2 mg/kg as used here) but not higher dose, Ztoau et al [53] observed a delayed decrease in CIN activity, i.e., when tested at 24 hours but not 2.5 hours. Additionally, in contrast to Witten et al [35], Ztoau et al [53] did not observe increased CIN firing in slices in response to amphetamine. Finally, in contrast to the predominance of studies showing increased ACh activity, Fleming et al [28] using fiber photometry report decreased ACh transients in response to psychostimulants, consistent with our data.

Although it is increasingly clear that striatal cholinergic interneurons and cholinergic signaling play a critical role in psychostimulant responses, detailed understanding of this role and underlying mechanism remains elusive; indeed, we cannot definitively conclude from extant literature even whether psychostimulants increase or decrease striatal cholinergic activity. The difficulty lies in experimental and methodological differences. Older microdialysis studies have slow time resolution and administer cholinesterase inhibitors to allow accumulation of ACh; however, that blockade of ACh clearance can have potentially profound effects on the microcircuitry and release of other neurotransmitters [55]. With electrophysiology or cFOS activation studies, caution is warranted in conflating CIN cell activity and acetylcholine release as modulation at the terminal can alter this relationship; indeed, that is the purpose of autoreceptors, including muscarinic autoreceptors on CINs. Thus, increased CIN activity could activate autoreceptors resulting in less acetylcholine release. Moreover, as the Guzman et al [52] study demonstrates, we should be equally cautious in equating observed behavioral effects that might correlate with CIN cell activity specifically to cholinergic release. As CINs release both glutamate and acetylcholine [56], the diminished cCPP observed by Witten et al [35] when inhibiting CINs could be mediated by decreased CIN glutamate release, not ACh release, which reconciles those results with the conflicting findings from Guzman et al [52] who found no effect on cCPP from entirely ablating striatal ACh.

In our study, we directly measure in real-time fluctuating extracellular ACh at sub-second temporal resolution without interfering with the signaling mechanisms we are measuring (i.e., without cholinesterase inhibitors or removing/replacing extracellular fluid). Unlike the prolonged increase we observe in basal extracellular dopamine (Fig 2a), we do not observe any similar slow timecourse changes in ACh in either direction. While contrary to prior microdialysis studies, this is more consistent with the pulsatile nature of striatal cholinergic signaling where alterations in ’cholinergic tone’ arise from change in frequency and magnitude of transient pulses of ACh that are rapidly cleared by acetylcholinesterase rather than changes in basal extracellular ACh [57, 58]. The overall decrease in ACh activity we observe could arise from decreased CIN activity, though unlikely as Witten et al [35] show increased activity and Ztoau et al [53] show no change in response to cocaine. Alternatively, it could arise from terminal modulation or altered synchrony in CIN firing [59, 60]. Notably, the Guzman finding that ablating striatal cholinergic signaling enhances behavioral sensitization is consistent with our observation of decreased ACh activity co-occurring with behavioral sensitization, though interestingly the sensitization of this decreased ACh activity itself is not necessary for behavioral sensitization. Finally, although a logical mechanism for decreased ACh activity in response to psychostimulants would be inhibition via D2R, our selective deletion of D2R in cholinergic neurons demonstrates that D2R is not mediating this decrease, only its sensitization (see below).

### 3.2 Dopamine-acetylcholine interaction

Considerable evidence indicates a robust phasic coupling between DA and ACh signals: reward-evoked DA transients are followed by brief pauses in CIN firing [61–63], coherent DA-ACh oscillations are observed in both dorsal [36] and ventral (our study) striatum, and the frequency of ACh transients predicts and controls cocaine extinction and conditioned place preference [28]. Therefore, a straightforward assumption would be that this interplay is created and maintained by direct receptor-level interactions—such as D2 or D5 receptors on CINs and nicotinic receptors on DA terminals. However, recent findings challenge this interpretation: Pharmacological blockade of DA or nicotinic receptors fails to disrupt DA-ACh coupling [36], suggesting that the coordinated transients arise largely from upstream glutamatergic afferents to CINs [30, 64] rather than reciprocal interactions between acetylcholine and dopamine systems themselves. Our results support this latter perspective: conditional deletion of D2Rs from CINs does not abolish the temporal relationship of their respective transients, indicating that a direct D2R mechanism– i.e., the dopamine to ACh linkage in reciprocal regulation– is not a fundamental driver of striatal DA-ACh co-fluctuations. We found that psychostimulants did not alter the temporal coordination of DA-ACh co-fluctuations, but did diminish the magnitude of their correlation/coherence. Both the drug and D2R KO data suggest that reciprocal interactions between DA and ACh within the microcircuitry of the striatum likely serves a modulatory role rather than driving the coordination between these two signaling systems.

### 3.3 Revisiting sensitization: dissociable components

Both KO and WT exhibit increased DA and decreased ACh. What differentiates the genotypes is how these signals adapt across repeated injections, where WT exhibit further progressive enhancement, i.e., sensitization, in both these responses across injections while the KO mice do not. This suggests that the primary responses and the sensitization of those responses are mediated by different mechanisms with the latter but not the former dependent upon CIN D2Rs.

The primary reduction in ACh in the absence of CIN D2Rs could arise from increased presynaptic inhibition of glutamate release driving CIN activity via D2R expressed on glutamatergic terminals and/or reduced synchrony between CINs [39, 65, 66], diminishing both the frequency and amplitude of transients. The lack of sensitization of this response in the absence of CIN D2Rs could point to D2Rmediated synaptic plasticity in CINs which might induce LTD in conjunction with reduced glutamatergic drive.

Loss of CIN D2Rs also abolishes sensitization in the DA response to psychostimulants and why this might be is not immediately clear. Intuitively, a reduction in Ach tone would be expected to decrease not increase facilitation of DA release via nicotinic receptors on dopamine terminals. Notably, D2R is deleted from all cholinergic cells, not just CINs. The most likely explanation is the loss of sensitization of increased DA release arises through the loss of D2Rs on different population of cholinergic neurons, such as those from the LTDg or PPTg to midbrain dopamine cells, again likely reflecting loss of D2R mediated adaptive plasticity.

Finally, it has long been thought that sensitization of DA mediates behavioral sensitization; however, our CIN D2R KO data suggests this is not correct. Despite complete loss of sensitization of the DA and ACh response to psychostimulants, behavioral sensitization is intact, but delayed, as observed both here and in Lewis et al [37]. This suggest that behavioral sensitization is driven by increased DA, perhaps mediating associative learning [67] but not its progressive enhancement. The role of the progressive enhancementof both DA and ACh responses to psychostimulants may be to accelerate behavioral sensitization rather than mediate it *per se*. This is particularly likely as the degree to which behavioral sensitization is slowed in the absence of CIN D2R appears to be dose dependent. We observed less delayed behavioral sensitization at 20 mg/kg than Lewis et al [37] observed at 10 mg/kg, suggesting that the magnitude of increased extracellular dopamine drives the rate of behavioral sensitization, consistent with the idea that sensitization of DA accelerates but does not mediate behavioral sensitization.

## Acknowledgments

This work was supported by NIH National Institute on Drug Abuse grant DA052871 (JAB), the Spar Biosciences Laboratory (JAB) generously funded by Dr. Ira Spar and by a PSCCUNY Award, jointly funded by The Professional Staff Congress and The City University of New York (JAB)

G.L. thanks Rudolf Faust for training and mentorship. G.L. also thanks Gaozhen Li, Jonathan Nudman, and Candice Gordon for their help and support in the lab. In addition, G.L. acknowledges Rudolf Faust, Youcef Bouchekioua, and Aske Lykke Ejdrup for their insightful discussions. We thank Paul Vezina for helpful input on designing the sensitization protocol.

## Author Contributions

G.L. constructed the experimental setup and performed the primary experiment (including virus injection and fiber optic placement surgery and subsequent photometry recordings) and F.G. conducted the experiment with D2R-CIN-KO mice. G.L. analyzed the data, including from the D2R-CIN-KO experiment, and wrote the manuscript. R.F. contributed to study design and trained GL. J.B. conceived the original idea, provided funding and laboratory resources and co-wrote the manuscript, providing feedback and guidance throughout the project.

## Methods

### 1 Animals

Sixteen wild-type animals were used for psychostimulant-induced locomotor sensitization experiments (8 male, 8 female, C57BL/6 Jackson Laboratory, 20-25 weeks old) and 9 mice (4 KO, 2 male, 5 WT, 3 male) were used for the D2R-CIN-KO experiment. The KO mice were generated by crossing mice carrying a conditional *Drd2* null allele (B6.129S4 (FVB)-Drd2^tm1.1Mrub^/J, JAX #020631; hereafter *fdrd2* ) bred to homozygosity for the *Drd2* null allele with mice expressing Cre-recombinase under the control of the choline acetylcholinesterase (ChAT) promoter (B6N.129S6(B6)Chat*^tm2(cre)Lowl^* /J, JAX #018957; hereafter *Chat-cre*). After crossing, mice heterozygous for the Cre-recombinase allele (Chat-cre^+/-^) that deleted D2R were used as experimental animals, while Cre-recombinase-null littermates (Chat-cre^-/-^) served as ‘wild-type’, non-D2R-deleted controls. All mice were age- and sex-matched with their respective control group. Animals were individually housed under a 12 h day/night cycle with *ad libitum* access to water and standard lab chow. All experiments were approved by the Queens College Institutional Animal Care and Use Committee.

#### 1.1 Stereotactic Surgery

Anesthesia was initiated with intraparetoneal (i.p.) ketamine/xylazine (90-120 mg/kg and 5-10 mg/kg, respectively) and maintained with 1-1.2 % isofluorane delivered through a nose cone. For photometry, we injected GRAB-rDA [68] and GRAB-ACh3.0 [69] with a titer of 2 ∗ 10^13^GC/ml and a volume of 175 nl per virus (350 nl per surgery) unilaterally (left hemisphere) into the nucleus accumbens shell at (AP 1.3 mm, ML -0.7 mm, DV 4.4 mm). The fiber optic implant was positioned at the site of virus injection and secured with metabond dental cement. Animals were allowed minimally four weeks for recovery and viral expression of sensors. We fabricated fiber optic implants in-house using 200 um fibers with a numerical apperture of 0.37 (Thorlabs, FT200UMT 0.39 NA, Ø200 µm Core Multimode Optical Fiber) and ceramic ferrules with dimensions of 1.25 mm x 6.4 mm and a bore size of 230 um (Thorlabs, CFLC230-10). The fiber was cut to the desired length and glued into the ferrule. Then, both ends of the fibers were polished until they had an efficiency of *>* 85%.

#### 1.2 Experimental Paradigm

Experiments were performed in the dark. Animals were habituated to an open-field arena (black spray-painted cylinder, 21 cm in diameter, 40 cm high), to the fiber-optic patch cable, and to (saline) IP injections over three days. To reduce stress during the IP injection, mice were habituated to handling for 2 weeks before the experiments. All sessions, including the habituation sessions, were 60 minutes with the IP injection being administered after the first 10 minutes. IP injections were performed in under one minute, and data collected (video, photometry) during that brief period were tagged and later removed from analysis.

Conventional video (infrared, ELP Camera USB 1080P), depth-camera (Microsoft Kinect 2) and fiber photometry data were recorded simultaneously. Photometry data were recorded as described below. A 3D-printed mount fixed the rotary joint above the chamber and kept the patch cable at optimal length. Video and photometry data streams were simultaneously collected and synchronized using Bonsai. Kinect data was synchronized with custom Python scripts.

Mice were tested for drug-induced locomotor sensitization via the following procedure:

After habituation, we acquired for every mouse a 60-minute baseline photometry recording with a saline injection after 10 minutes. After three days without injections, we recorded a first injection of either cocaine (20 mg/kg, n = 5), d-amphetamine (2 mg/kg, n = 5), or saline (n=6) via the same protocol, i.e., after 10 minutes pre-injection baseline recording. We administered five additional injections over 5 consecutive days, acquiring a second drug-session photometry recording at injection/day 5. Animals were then given 7 days without any activity or injection before all animals were administered a challenge injection, to compare differences in response between drug-naive animals (the saline group) and animals previously exposed cocaine and amphetamine. Previously exposed animals were administered the same drug and drug-naive (saline group) mice were administered either cocaine (n = 3) or d-amphetamine (n = 3) at the same doses. In the D2R-CIN-KO study, we did not have a saline or drug-naive group but in describing the results continued to use the term ’challenge’ dose for consistency with regard to the experimental timeline and procedures.

### 2 Behavioral Analyses

#### 2.1 DeepLabCut

We utilized DeepLabCut [38], an open-source, markerless pose estimation software, to track specific body parts of mice in video recordings. The model was fine-tuned on a pre-trained ResNet-50 neural network initialized with ImageNet weights. Video data were captured using an infrared camera at high-definition resolution and downsampled to a height of 216 pixels to optimize computational efficiency.

A total of 269 frames were manually annotated, labeling all visible instances of the following body parts: nose, left ear, right ear, left eye, right eye, four segments of the tail (*tail 1* to *tail 4* ), left forelimb, right forelimb, left backlimb, and right backlimb. Only body parts visible in each frame were annotated to ensure accurate training data. The model was trained using data augmentation techniques to improve generalization. Training proceeded for 450,000 iterations, with the checkpoint exhibiting the lowest loss at 135,000 iterations selected for subsequent analysis. This model achieved a training error of 2.72 pixels and a test error of 2.83 pixels, measured as the average Euclidean distance between predicted and annotated positions. Applying a confidence cutoff (*pcutoff* ) of 0.6 improved the training error to 1.92 pixels and the test error to 2.67 pixels.

Post-training, all videos were processed through the trained model to perform pose estimation. We used a confidence threshold of *p* = 0.6 for all labels to filter out low-confidence predictions. The filterpredictions() function in DeepLabCut was applied to smooth the output trajectories and reduce potential noise. Unless otherwise specified, default parameters were used throughout the process.

#### 2.2 Velocity and Distance

We quantified locomotor activity by tracking the base of the tail in video recordings using DeepLabCut (see 2.1). The pixel coordinates obtained were converted to centimeters using a scaling factor from a previous calibration. The distance between each consecutive frame was calculated to determine movement over time. Total distance traveled during each session was obtained by summing these frame-to-frame distances. For temporal analysis, data were binned into 1-minute intervals. The average velocity for each bin was calculated by summing the distances within that minute and dividing by 60 seconds, allowing us to assess changes in locomotor activity over time.

#### 2.3 Time without Locomotion and Center Time

To quantify periods without locomotion, we implemented a rest detection algorithm with two key parameters: a minimum duration threshold of 2 seconds and a movement threshold of 1 pixel between consecutive frames. A rest period was defined when frameto-frame movement remained below this threshold for at least 2 seconds. The total time without locomotion was calculated by summing all identified rest periods and converted to minutes.

Center time was quantified using an automated center point detection algorithm. The arena center was determined by fitting a circle to the mouse’s tracked positions throughout the session using least squares optimization. For each frame, we calculated the distance between the mouse’s nose position and the computed arena center, converting these distances from pixels to centimeters using a calibration factor. Time in center was defined as periods when the mouse’s distance to center was less than 5 cm. The total center time was calculated by summing all frames meeting this criterion.

### 3 Fiber Photometry

#### 3.1 Data Acquisition

All photometry recordings were performed with a commercial photometry system by Neurophotometrics (FP3002) [70]. Three LEDs at frequencies of 560 nm (red-shifted indicators for dopamine), 470 nm (green indicators for acetylcholine), and 410 nm (isosbestic control signal) were used. Only one LED was on at a time and recordings were performed at a frequency of 90 Hz, which effectively yields 30 Hz. The system was connected to fiber optic cables attaching to the mice via a low-autofluorescence patch cord (Doric Lenses, Quebec, Ontario) using a pigtailed rotary joint (Doric Lenses). We used low power excitation light at 50 uW.

#### 3.2 Data Analysis

Raw photometry data were processed using custom Python scripts using standard scientific computing libraries (numpy, pandas, scipy). The preprocessing pipeline included several key steps:

1. Frame drops were detected and quantified based on Neurophotometric’s frame counter timestamps. All sessions showed minimal frame drops (*<* 1% of total frames), with most sessions showing no frame drops.
2. Signal alignment was performed by shifting reference traces (isosbestic control) by 1-2 frames relative to the sensor signals to account for the sequential LED illumination pattern. The specific shift was determined based on the recording channel (470 nm or 560 nm).
3. Recording segments were temporally aligned to behavioral data using computer timestamps that simultaneously logged both photometry and behavioral (video) data. The experimenter additionally marked their presence in the recording room for IP-injection by pressing a keyboard button upon entry and exit. These timestamps were used to confirm that all interventions occurred within a one-minute window that was subsequently excluded from analysis.
4. During IP-injections, recordings continued uninterrupted and the optical fiber remained untouched to maintain a stable baseline. Visual inspection of the isosbestic control signal confirmed the absence of baseline shifts during the injection period. Each session included 10 minutes of pre-injection baseline, one minute for IP-injection, and 50 minutes of post-injection recording, totaling 61 minutes of data.

#### 3.3 Detrending of Dopamine Traces

To detrend baseline sessions, a bi-exponential curve *F_fit_* was fitted to each recording using non-linear least squares:

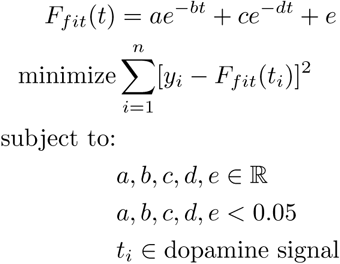

This function models the gradual decay in fluorescence signal due to photobleaching as the sum of two exponential decay processes plus an offset. Where:

- *F_fit_*(*t*) is the biexponential function modeling photobleaching
- *a, c* are amplitude parameters for the two exponential components
- *b, d* are decay rate parameters
- *e* is the offset parameter
- *y_i_* are the observed fluorescence values
- *t_i_* are the timepoints during baseline period
- The parameters are constrained to be less than 0.05 to avoid physiologically implausible values

The fluorescence signal was converted to Δ*F/F* by subtracting and dividing by the fitted photobleaching function:

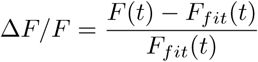

where *F* (*t*) is the raw fluorescence signal and *F_fit_*(*t*) is the biexponential fit modeling photobleaching: *F_fit_*(*t*) = *ae^−bt^* + *ce^−dt^* + *e*. This normalization corrects for photobleaching while preserving relative changes in fluorescence that reflect neural activity. Z-scores (*z*Δ*F/F* ) where calculated by standardizing to the mean and the standard deviation of the Δ*F/F* during the 10-minute pre-injection period.

Artifacts created by the rotary joint were removed by fitting and subtracting a baseline with the AirPLS algorithm to the isosbestic and the DA signal, scaling the isosbestic using non-negative robust linear regression (sklearn.linear model.Lasso), and subtracting the scaled isosbestic fit from the DA photometry signal.

For injection 1, 5, and challenge, post-injection data weren’t used to fit the biexponential decay and for all recordings, only pre-injection data were used to calculate mean and std for standardization (z-scoring).

#### 3.4 DA Transients

Dopamine transients were identified using a peak detection algorithm on the z-scored Δ*F/F* traces. The signal was first smoothed using a 5-point centered rolling average to reduce random noise. Local maxima and minima were detected by finding zero-crossings in the first derivative of the smoothed signal. A transient was defined as an event where the amplitude difference between a local minimum and the preceding local maximum exceeded a threshold of 1 z-score. The amplitude of each detected transient was calculated as the difference between the local maximum and minimum values. This approach allowed for automated detection of rapid increases in dopamine signaling while being robust against slower baseline fluctuations.

#### 3.5 Decay of DA Transients

To quantify the decay kinetics of dopamine transients, we fit an exponential decay function to the falling phase of each detected transient. First, transients were identified as outlined above 3.4. For each transient, we analyzed the time window from 0.2 to 5 seconds after the peak to focus on the decay phase while avoiding the initial rise. The decay was modeled using a single exponential function of the form *f* (*t*) = *a* · *e^−t/τ^* , where *τ* represents the decay time constant. Curve fitting was performed using scipy’s curve fit function with initial parameter estimates of *a* = 1 and *τ* = 0.1*s*. The half-life (*t*_1*/*2_) was calculated from the fitted *τ* as *t*_1*/*2_ = ln(2) · *τ*. This analysis was performed separately for transients detected during different experimental phases (baseline, rise and plateau) to examine how drug treatments affected dopamine clearance kinetics.

#### 3.6 Wavelet Transform

To analyze the frequency components of the dopamine signal, we performed a discrete wavelet transform using the Symlet-4 (sym4) wavelet. The dopamine signal (Δ*F/F* ) was decomposed into different frequency bands using the `pywt.wavedec` function from the PyWavelets library, with the maximum decomposition level determined by the signal length. For each frequency band, we calculated the amplitude as the square root of the sum of squared wavelet coefficients. To compare the frequency components between experimental groups, we normalized the amplitudes to those measured in receptor-mutant mice (rDA-mut) under vehicle conditions, computing a fold-change for each frequency band. Statistical analysis was performed using a linear mixed-effects model with substance and frequency band as fixed effects and mouse ID as a random effect, allowing for interaction between substance and frequency band (`fold change` ∼ `Substance * Frequency Band`). The resulting frequency bands ranged from ultraslow (9 min periods) to fast oscillations (∼ 15 Hz), with the highest possible frequency being the Nyquist frequency (sampling rate/2 = 15 Hz).

#### 3.7 ACh Transients

Acetylcholine transients were detected using scipy’s `find_peaks` algorithm. Peaks were identified as local maxima in the z-scored fluorescence trace (*z*Δ*F/F* ) that exceeded a threshold of 2 z-scores above baseline. To prevent the duplicate detection of spurious peaks due to noise, we implemented a minimum horizontal distance criterion of 100 ms between successive peaks. This approach allowed for the automated detection of discrete acetylcholine release events while minimizing false positives from high-frequency noise in the signal.

#### 3.8 Cross-Correlation Analysis

To quantify transient-scale coordination between DA and ACh signals, we computed cross-correlations between the detrended DA signal and ACh in 2-second windows. This procedure isolates moment-to-moment interactions by minimizing the confounding influence of long-lasting drug-induced changes. The significance of cross-correlations was assessed by comparing observed values to shuffled distributions.

#### 3.9 Coherence Analysis

We computed the coherence between DA and ACh signals (fs=30 Hz) to identify frequency bands where the signals co-fluctuate. Coherence was estimated using standard spectral analysis techniques (e.g., Welch’s method or related approaches). To determine significance, coherence values were compared against surrogate distributions generated by circularly shifting one signal relative to the other. Changes in coherence before and after drug injections were tested using linear mixed-effects models.

#### 3.10 Phase Offset

Phase offsets between ACh and DA were computed using cross-spectral density methods. Because we were interested in the stability of the temporal relationship, we focused on the average phase offset near the primary coherence band ( 0.5–1 Hz). Phase offsets were then compared across groups and sessions to test for drug-induced shifts.

#### 3.11 Event-Triggered Analysis

We identified DA transients (e.g., those exceeding 1 z-score thresholds) and aligned ACh signals to these event times. For each DA event, we measured ACh activity in a window of several seconds before and after the DA peak. By averaging across many events, we obtained a time-resolved profile of how ACh changes around DA transients. This analysis reveals whether ACh fluctuations systematically precede or follow DA events and how this relationship is affected by psychostimulants.

### 4 Statistical Tests

We used a mixed-effects linear regression model (LME) to evaluate the effects of *Group* (saline, cocaine, amphetamine), *Session* (baseline, injection 1, injection 5, challenge), or Genotype (KO, WT), and their interaction on neural and behavioral measurements. The model included fixed effects for *Group* (or *Genotype*), *Session*, and their interaction, and random effects for each mouse to account for individual variability:

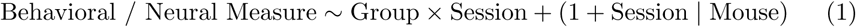

where Mouse is the grouping variable for random intercepts and slopes with respect to *Session*. Statistical significance was assessed using *z* -tests on the estimated coefficients, and *p*-values were reported with a significance threshold of 0.05.

## Supplemental Material

**Fig. 7:**
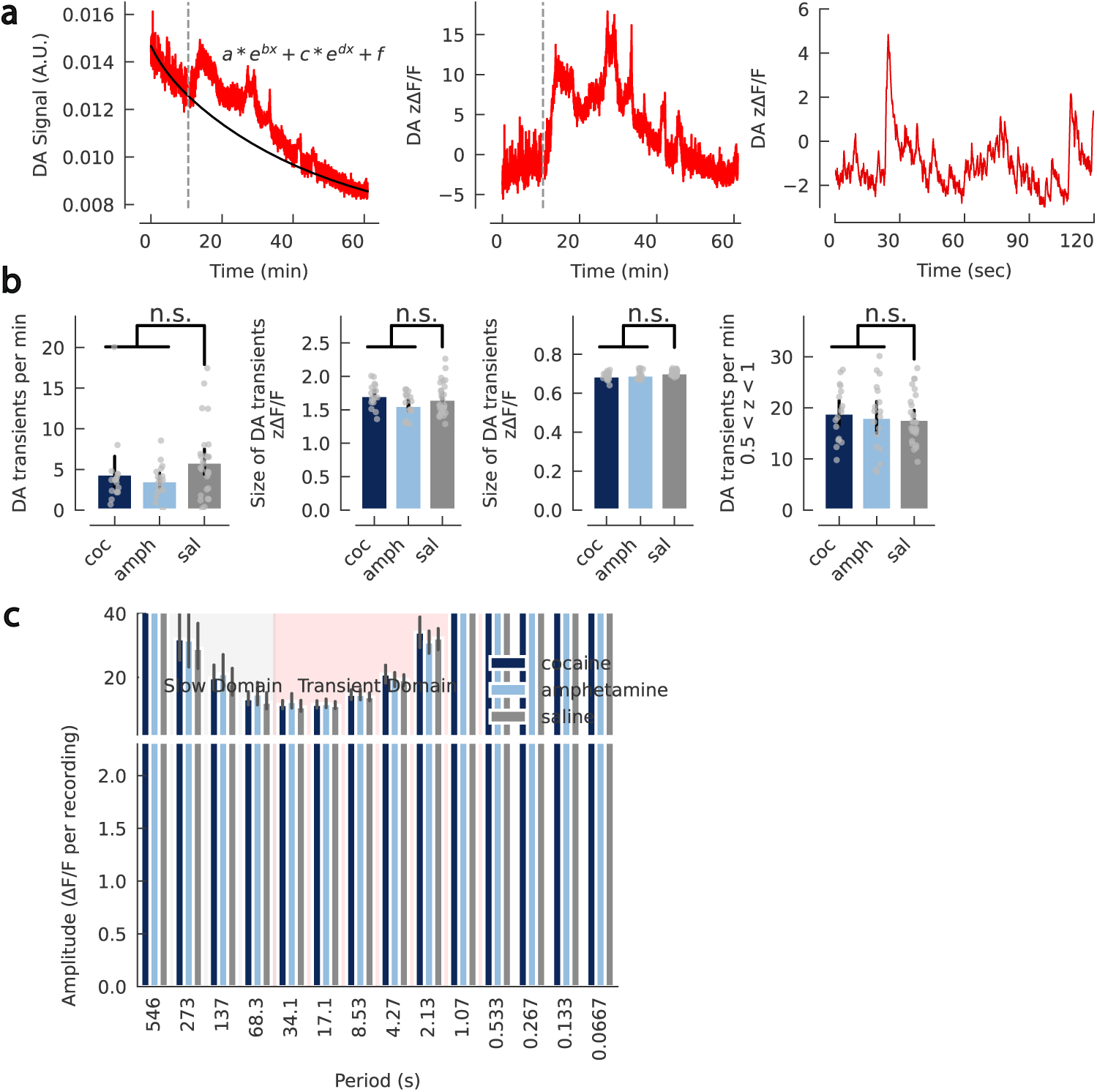
Supplementary Figure: Dopamine signal preprocessing and transient analysis. **a**, Left panel: raw photometry data from one example session (red) with fitted baseline (black). Middle panel: detrended and normalized dopamine fluorescence (zdFF) for the same session. Right panel: zoom-in of the middle panel highlighting dopamine transients. **b**, Left panel: pre-injection controls for the number and size of dopamine transients. No significant differences were observed in the number of transients compared to saline controls (cocaine: *p* = 0.13; amphetamine: *p* = 0.33) or in their size (cocaine: *p* = 0.84; amphetamine: *p* = 0.13). Right panel: number and size of dopamine peaks with amplitudes between 0.5 and 1 z-scores. The number of such peaks showed a trend toward significance for cocaine (*p* = 0.065) but not for amphetamine (*p* = 0.70). Their size was significantly different for cocaine (*p* = 0.013) and showed a trend for amphetamine (*p* = 0.055). **c**, Dopamine amplitude per frequency obtained via wavelet analysis. Error bars represent the 95% bootstrap confidence interval. Statistical significance was assessed using linear mixed-effects models with appropriate fixed and random effects. Significance stars are indicated as \**p <* 0.05, \*\**p <* 0.01, \*\*\**p <* 0.001.

**Fig. 8:**
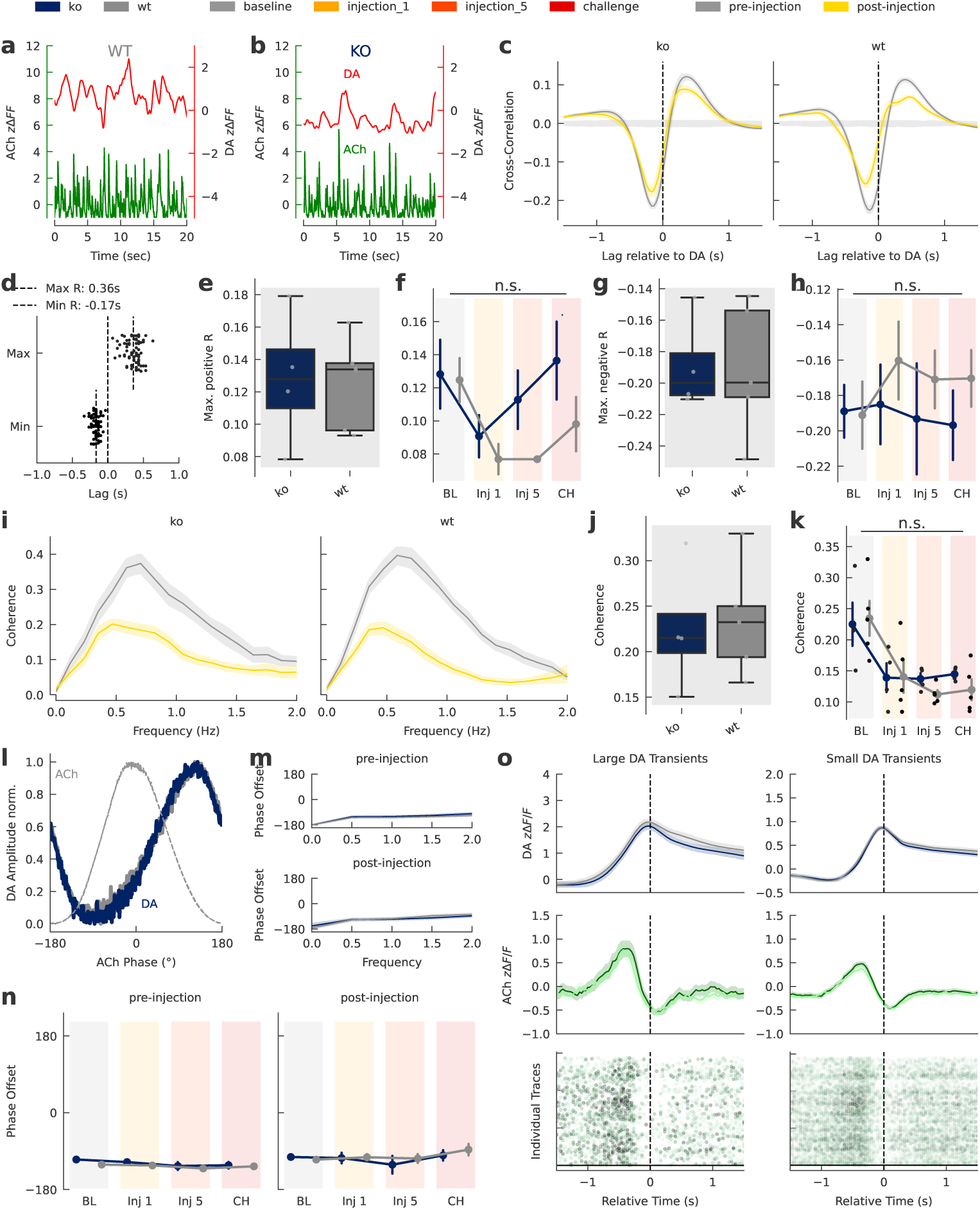
Conditional D2R KO in CINs does not change DA–ACh interaction. **a**, A 20-s DA (red) and ACh (green) trace from a representative WT animal. **b**, DA and ACh from a KO animal. **c**, Cross-correlation between DA and ACh fluorescence before and after IP injection for KO (left) and WT (right) mice; lag is relative to DA. **d**, Time of the largest positive and negative crosscorrelation post-injection. Each dot represents the average of one recording. **e**, Max. positive Pearson’s R between DA and ACh fluorescence at injection 1 for KO and WT animals (LME, *p* = 0.19). **f**, Max. positive Pearson’s R by genotype and session (LME, Genotype*×*Session interaction: *p* = 0.24). **g**, Max. negative Pearson’s R between DA and ACh fluorescence at injection 1 (LME, *p* = 0.28). **h**, Max. negative Pearson’s R by genotype and session (LME, Genotype*×*Session interaction: *p* = 0.86). **i**, Coherence before and after cocaine injection for KO (left) and WT (right) mice. **j**, Mean coherence (0–1 Hz) between DA and ACh fluorescence at injection 1 (LME, *p* = 0.63). **k**, Mean coherence (0–1 Hz) by genotype and session (LME, Genotype*×*Session interaction: *p* = 0.79). **l**, Mean normalized DA fluorescence versus the phase of periodic ACh fluctuations (grey for reference). **m**, Phase offset of DA versus ACh fluorescence by genotype. **n**, Phase offset (degrees) between DA and ACh for the pre-injection period (LME, no significant effects) and the post-injection period (LME, no significant effects). **o**, Upper panel: DA fluorescence time-locked to the peak of detected transients per genotype. Middle panel: mean ACh fluorescence during the same events. Bottom panel: individually detected ACh events around DA transients, colored by height (dashed lines separate animals). Left panels show data for DA transients larger than 2 z-scores; right panels, 1–2 z-scores. Error bands: s.e.m. (line and point plots), 95% bootstrap CI (bar plots). Significance: \**p <* 0.05, \*\**p <* 0.01, \*\*\**p <* 0.001.

