## supplemental material for "Accumbal Dopamine and Acetylcholine Dynamics during Psychostimulant Sensitization"

### Appendix E Supplemental Material

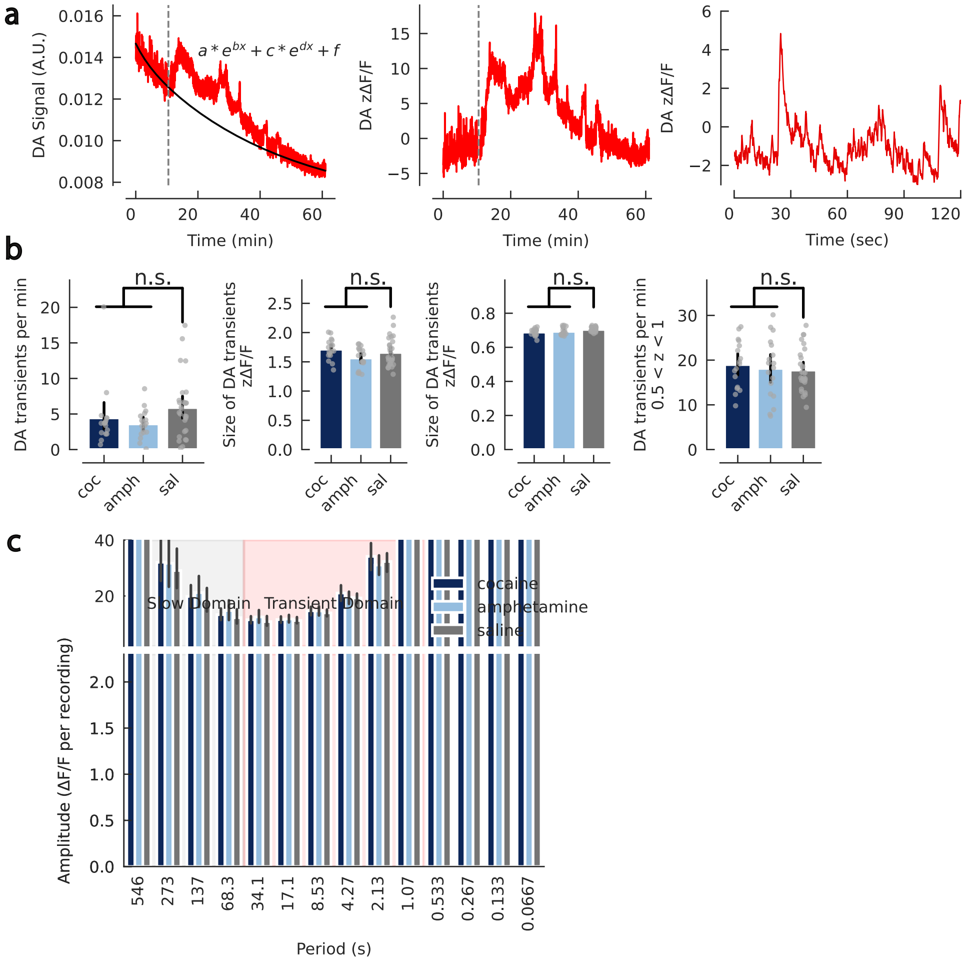


**Fig. E1**: **Supplementary Figure: Dopamine signal preprocessing and transient analysis.** **a**, Left panel: raw photometry data from one example session (red) with fitted baseline (black). Middle panel: detrended and normalized dopamine fluorescence (zdFF) for the same session. Right panel: zoom-in of the middle panel highlighting dopamine transients. **b**, Left panel: pre-injection controls for the number and size of dopamine transients. No significant differences were observed in the number of transients compared to saline controls (cocaine: $p = 0.13$; amphetamine: $p = 0.33$) or in their size (cocaine: $p = 0.84$; amphetamine: $p = 0.13$). Right panel: number and size of dopamine peaks with amplitudes between 0.5 and 1 z-scores. The number of such peaks showed a trend toward significance for cocaine ($p = 0.065$) but not for amphetamine ($p = 0.70$). Their size was significantly different for cocaine ($p = 0.013$) and showed a trend for amphetamine ($p = 0.055$). **c**, Dopamine amplitude per frequency obtained via wavelet analysis. Error bars represent the 95% bootstrap confidence interval. Statistical significance was assessed using linear mixed-effects models with appropriate fixed and random effects. Significance stars are indicated as $^{*}p < 0.05$, $^{**}p < 0.01$, $^{***}p < 0.001$.

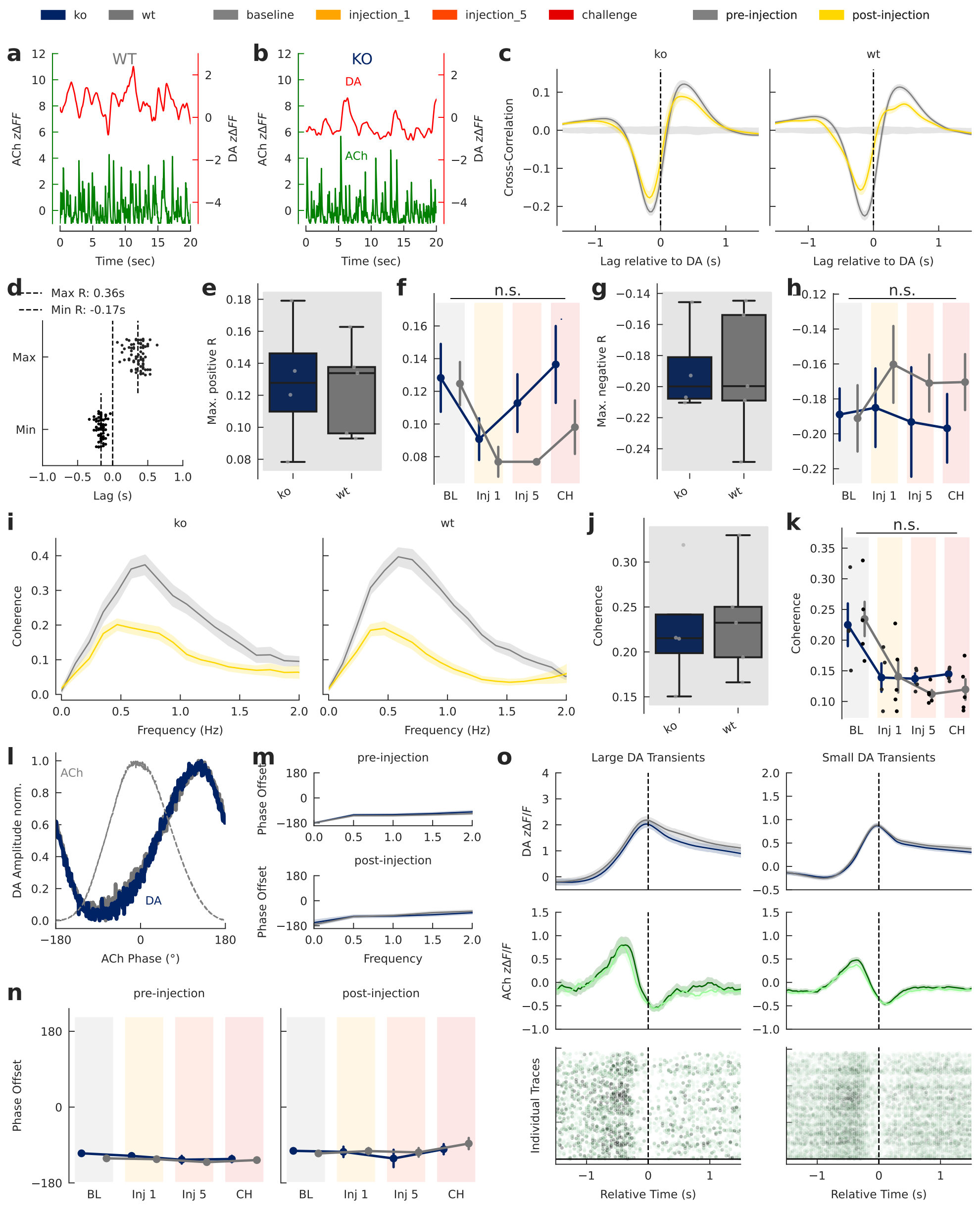


**Fig. E2**: **Conditional D2R KO in CINs does not change DA–ACh interaction.** **a**, A 20-s DA (red) and ACh (green) trace from a representative WT animal. **b**, DA and ACh from a KO animal. **c**, Cross-correlation between DA and ACh fluorescence before and after IP injection for KO (left) and WT (right) mice; lag is relative to DA. **d**, Time of the largest positive and negative cross-correlation post-injection. Each dot represents the average of one recording. **e**, Max. positive Pearson's R between DA and ACh fluorescence at injection 1 for KO and WT animals (LME, $p = 0.19$). **f**, Max. positive Pearson's R by genotype and session (LME, Genotype×Session interaction: $p = 0.24$). **g**, Max. negative Pearson's R between DA and ACh fluorescence at injection 1 (LME, $p = 0.28$). **h**, Max. negative Pearson's R by genotype and session (LME, Genotype×Session interaction: $p = 0.86$). **(cont.)**

**Fig. E2**: **(cont.) Conditional D2R KO in CINs does not change DA–ACh interaction.** **i**, Coherence before and after cocaine injection for KO (left) and WT (right) mice. **j**, Mean coherence (0–1 Hz) between DA and ACh fluorescence at injection 1 (LME, $p = 0.63$). **k**, Mean coherence (0–1 Hz) by genotype and session (LME, Genotype×Session interaction: $p = 0.79$). **l**, Mean normalized DA fluorescence versus the phase of periodic ACh fluctuations (grey for reference). **m**, Phase offset of DA versus ACh fluorescence by genotype. **n**, Phase offset (degrees) between DA and ACh for the pre-injection period (LME, no significant effects) and the post-injection period (LME, no significant effects). **o**, Upper panel: DA fluorescence time-locked to the peak of detected transients per genotype. Middle panel: mean ACh fluorescence during the same events. Bottom panel: individually detected ACh events around DA transients, colored by height (dashed lines separate animals). Left panels show data for DA transients larger than 2 z-scores; right panels, 1–2 z-scores.
Error bands: s.e.m. (line and point plots), 95% bootstrap CI (bar plots). Significance: $^{*}p < 0.05$, $^{**}p < 0.01$, $^{***}p < 0.001$.
